# *Xanthomonas* transcriptome inside cauliflower hydathodes reveals bacterial virulence strategies and physiological adaptation at early infection stages

**DOI:** 10.1101/2021.01.19.427230

**Authors:** Julien S. Luneau, Aude Cerutti, Brice Roux, Sébastien Carrère, Marie-Françoise Jardinaud, Antoine Gaillac, Carine Gris, Emmanuelle Lauber, Richard Berthomé, Matthieu Arlat, Alice Boulanger, Laurent D. Noël

**Author notes:** Authors for correspondence and, LIPME, Université de Toulouse, INRAE, CNRS, Université Paul Sabatier, Castanet-Tolosan, France. These authors contributed equally to this work (Julien S. Luneau, Aude Cerutti).

## Abstract

*Xanthomonas campestris* pv. *campestris* (*Xcc*) bacterium is a seed-transmitted vascular pathogen causing black rot disease on cultivated and wild *Brassicaceae. Xcc* enters the plant tissues preferentially via hydathodes which are organs localized at leaf margins. In order to decipher both physiological and virulence strategies deployed by *Xcc* during early stages of infection, the transcriptomic profile of *Xcc* was analyzed three days after entry into cauliflower hydathodes. Despite the absence of visible plant tissue alterations and a bacterial biotrophic lifestyle, 18% of *Xcc* genes undergo a transcriptional reprogramming, including a striking repression of chemotaxis and motility functions. *Xcc* full repertoire of virulence factors was not yet activated but the expression of the 95-gene HrpG regulon, including genes coding for the type three secretion machinery important for suppression of plant immunity, was induced. The expression of genes involved in metabolic adaptations such as catabolism of plant compounds, transport functions, sulfur and phosphate metabolism was upregulated while limited stress responses were observed three days post infection. These transcriptomic observations give information about the nutritional and stress status of bacteria during the early biotrophic infection stages and help to decipher the adaptive strategy of *Xcc* to the hydathode environment.

## Introduction

*Xanthomonas campestris* pv. *campestris* (*Xcc*) is a seed-transmitted vascular pathogen causing black rot disease on *Brassicaceae*. This bacterium has a complex lifestyle composed of both epiphytic and endophytic stages (1) which have been studied using molecular genetics since the 80’s (2). *Xcc* epiphytic life is associated with environmental stresses such as UV or dehydration and relies, for instance, on the production of xanthan exopolysaccharides (EPS) or protective pigments such as xanthomonadin. Upon favorable conditions, *Xcc* will gain access into the leaf inner tissues via wounds or hydathodes (3).

Hydathodes are plant organs localized at the leaf margin mediating guttation. Hydathodes are classically composed of an epidermis with water pores resembling stomata and an inner lose parenchyma called epithem irrigated by numerous xylem vessels (4). These specific structures offer an ecological niche for pathogenic bacteria and a rapid access to xylem vessels leading to systemic vascular infections (3). However, only few pathogens have been demonstrated to colonize this niche and the conditions driving hydathode infection are poorly understood (3, 5-9). While a pre-invasive immunity limiting *Xcc* entry through water pores could not be evidenced, a post-invasive immunity was described inside the epithem (3). This is best revealed by the inability of a bacterial mutant of the Hrp (hypersentive response and pathogenicity) type III secretion (T3S) system to multiply in the epithem and to initiate vascular infections. The T3S system is responsible for the secretion and translocation of type III effector (T3E) proteins inside plant cells where they interfere with plant physiology and suppress plant immunity (10). These results highlight the importance of immune suppression for the establishment of the infection.

Once inside hydathodes, *Xcc* adapts to this niche and adopts a biotrophic lifestyle: bacteria slowly multiply in the apoplastic spaces between epithem cells without causing visible tissue alterations as observed until three days post infection (dpi) of cauliflower hydathodes (3). A switch to a necrotrophic behaviour is then observed, resulting in the almost complete digestion of the epithem at 6 dpi and *Xcc* vascularization. Systemic infection reaching the flowers will cause seed colonization and transmission to seedlings (11, 12).

During infection, *Xcc* may feed on guttation fluid, xylem sap or plant tissues (13). This process can be facilitated by high affinity nutrient transport systems such as TonB-dependent transporters (TBDT) for mineral (e.g iron) or carbohydrate nutrition (e.g. sucrose, 14) and to a large repertoire of plant cell wall degrading enzymes secreted through the type two secretion (T2S) system. Such metabolic adaptations need to be finely coordinated throughout the infectious cycle. Master regulators include the *rpf*-DSF (Diffusible Signal Factor) quorum sensing system, sensors of nutrient availability and metabolic activity and two-component systems that allow bacterial cells to respond appropriately to diverse extracellular stimuli encountered during its life cycle such as oxidative stress, oxygen levels, pH, temperature and plant signals (15-17). Among them, some are well known to play a major role in virulence such as the response regulator HrpG which is, with the transcription regulator HrpX, the master regulator of the T3S regulon (18-20).

Our knowledge of *Xcc* gene expression *in planta* remains elusive and technically challenging especially at early steps of infection when bacterial populations are low. Indeed, most transcriptomic studies in *Xcc* or other *Xanthomonas* species were performed so far *in vitro* focusing on specific regulons of the *hrpG* and *hrpX* genes in *hrp-inducing* media, of the *prc* protease gene, of the DSF-mediated quorum sensing system or of the *gum* genes responsible for xanthan production (21-31). Only few *in planta* transcriptomic analyses were performed on *Xanthomonas* (28, 32-34) and all at late stages of infection and/or in comparison to *in vitro*-grown bacteria. Such *in planta* transcriptomics approaches would help identify new pathogenic behaviors and adaptation to the host during the infection process and in the different tissues colonized. Such approaches are also good descriptors of the environmental conditions and stresses imposed by the host to the bacterial pathogen (35).

In this study, we compare the transcriptome of *Xcc* inside cauliflower hydathodes at 4 or 72 hours after inoculation, *i*.*e*. during the biotrophic stage of infection, in order to determine *Xcc* adaptative transcriptomic responses and to infer the environmental conditions met by this bacterial pathogen inside these plant organs.

## Materials and methods

### Bacterial strains, plasmids and growth conditions

The list of strains and plasmids used in this study is provided in Table S1. *Xcc* was cultivated in MOKA medium (4 g.l^-1^ Yeast extract, 8 g.l^-1^ Casamino acids, 1 mM MgSO_4_ and 2 g.l^-1^ K_2_HPO_4_) at 28°C under agitation at 200 rpm or on MOKA-agar plates (14). *E. coli* strains TG1 and strain carrying pRK2073 helper plasmid were cultivated in liquid LB medium or on LB-agar plates at 37°C under agitation. Antibiotics were used at the following concentrations: 50 µg/ml rifampicin, 50 µg/ml kanamycin, and 40 µg/ml spectinomycin.

### Mutagenesis and complementations

In-frame deletion mutants of *Xcc* were obtained by double recombination with derivatives of the suicide plasmid pK18mobSacB as described (36). Sequences flanking the deletion were amplified from *Xcc* strain 8004 genomic DNA and introduced into pK18mobSacB by Gibson assembly (37). For complementation, the CDS was amplified from *Xcc* strain 8004 genomic DNA and cloned by Gibson assembly into plasmid pK18_CompR3, a pK18mobSacB derivative containing a pTac promoter and a T7 terminator region from pCZ1016 (38) flanked by *XC_1301* and *XC_1302* sequences to drive stable insertion at the *XC_1301*/*XC_1302* interval. For genes lacking ribosome-binding site (RBS) in their upstream region, the RBS from plasmid pK18-GUS-GFP (3) was inserted downstream the pTac promoter giving pK18_compR3_RBS. All plasmids were conjugated into *Xcc* 8004::GUS-GFP derivatives by triparental mating with the *E. coli* TG1 carrying pRK2073 helper plasmid as described (3, 39, 40). The sequences of oligonucleotides used to construct deletion and complementation plasmids are listed in Table S2. The growth of all strains was assessed in MME and MOKA media (Figure S1).

### Plant growth conditions

*Brassica oleracea* var *botrytis* cv. Clovis F1 (cauliflower) were grown under greenhouse conditions. Four-weeks-old plants were transferred one day before inoculations in a growth chamber (9 hours light; 22°C; 70% relative humidity).

### Preparation of biological samples used for RNA sequencing

*Xcc* strains were grown *in vitro* in MOKA medium to mid-exponential phase, harvested by filtration as described (18) and stored at −80°C.

Hydathodes from the first three cauliflower leaves were inoculated by continuous or transient dipping as described (3) using 0.01% of SILWET-L77® (DE SANGOSSE). For continuous dipping, leaves were dipped in a bacterial suspension for 4 hours in a covered small greenhouse and immediately harvested. For transient dipping, leaves were dipped in a bacterial suspension for ca. 15 seconds, watered, placed in a small greenhouse, covered for a day and a half and harvested 72 hours post inoculation. To collect hydathodes, leaves were briefly rinsed twice in sterile distilled water and dried on a paper towel prior to harvesting. At least 1000 hydathodes were macrodissected per condition with a 1.5 mm diameter punch.

Collected tissues were immediately placed in RNA Protect Bacteria Reageant® (Quiagen™, 2:1 (v/v) of RNA Protect and RNAse free water). After three minutes sonication in a water bath, the supernatant was recovered and centrifuged for 10 minutes at 5000g. The pellet was stored at −80°C. Three independent biological replicates were obtained.

To determine the infection level of hydathodes under the two inoculation protocols used, the bacterial populations were determined at 4 and 72 hpi in 24 and 30 individual hydathodes respectively as described below (Figure S2).

### RNA extractions, ribodepletion and sequencing

RNA extraction and ribodepletion were performed as previously described (41). Oligonucleotide probes used for RNA depletion were directed against *Xanthomonas* rRNA and 2 tRNA (Ile and Ala) (18) and for plant-derived samples probes targeting Arabidopsis and *Brassica oleracea* rRNA and major chloroplastic RNA (41) (Table S3). RNAs were fractionated into short (<200 nt) and long (>200 nt) RNA fractions using Zymo Research RNA Clean & ConcentratorTM-5 columns (Proteigene) and subjected to oriented sequencing (See supplemental material for detailed procedures). Raw sequence data were submitted to the Sequence Read Archive (SRA) database (Accession SRP280320 and SRP280329).

### Reannotation of *Xcc* strain 8004 genome sequence

Annotation of *X. campestris* pv. *campestris* strain 8004 genome was performed using EuGene-PP (41) (EuGene-PP v1.0, eugene-4.1c) with SRP280320 RNA libraries and *X. campestris* pv. *campestris* strains 8004, ATCC33913 and B100 public annotations GCA_000012105.1, GCA_000007145.1 and GCA_000070605.1, respectively. This new annotation is available at https://dx.doi.org/10.25794/reference/id52ofys.

### Analysis of RNA sequencing results and statistical analysis

Mapping of RNA sequencing reads was performed on the *Xcc* strain 8004 reannotated genome sequence (42, Genbank accession number CP000050.1) and when appropriate on sequences of *Brassica oleracea* nuclear genome (Brassica_oleracea.v2.1.31; Accession GCA_000695525.1), *Brassica oleracea* mitochondrial genome (accession NC_016118.1) and *Brassica rapa* chloroplastic genome (BRARA_CHL, accession NC_040849.1) as described (18).

Differentially expressed genes (DEG) were detected with EdgeR Bioconductor package version 3.30.3 (43). Genes with no counts across all libraries were discarded. Normalization was performed using TMM (trimmed mean of M-values) method (44). Quality control plots of normalized data sets and reproducibility of biological repeats were generated by principal component analysis using Ade4 version1.7-15 package (45) and heatmaps obtained with the package pheatmap version 1.0.12 (Raivo Kolde (2015). pheatmap: Pretty Heatmaps. R package version 1.0.8. https://CRAN.R-project.org/package=pheatmap) on sample-to-sample Euclidean distances.

Fitted generalized linear models (GLM) with a design matrix Multiple factor (biological repetition and factor of interest) were designed. The Cox-Reid profile-adjusted likelihood (CR) method in estimating dispersions was used. DEG were called using the GLM likelihood ratio test using a False Discovery Rate (FDR) (46) adjusted *q*-value < 0.05. Clustering on filtered DEG (*q*-value < 0.05 in at least on biological condition) was generated with heatmap.2 function as available in the gplots Bioconductor package version 3.0.1. (47) using Ward’s minimum variance clustering method on Euclidean (48). Analysis of gene ontology enrichment was conducted using the topGO package version 2.40.0 (49).

### Infection of hydathodes and measurement of bacterial population

For hydathode infection, the second true leaf of cauliflower plants was dip-inoculated in a bacterial suspension at 10^8^ cfu/mL in 1mM MgCl_2_ containing 0.5% (v/v) Tween 80. In order to determine *Xcc* populations in single hydathodes, hydathodes were collected at three or six days post-inoculation by macrodissection with a 1.5 mm-diameter punch. Eight hydathodes were sampled per leaf and individually placed in 200 µL of 1mM MgCl_2_. After bead-assisted grinding at 30 Hz for 2 min using a Retsch MM400 grinder, 5-µL droplets of serial dilutions were spotted on MOKA plates supplemented with 30 µg/mL pimaricin in three technical replicates and incubated at 28°C for two to three days. Individual colonies were counted and the mean of the three technical replicates was calculated to estimate the infection level of each hydathode. Experiments were performed on three plants per condition and in three independent biological replicates.

Significance of differences observed in bacterial population quantifications and bacterial pathogenicity assays was assessed using the non-parametric Kruskal-Wallis test with α = 0.05.

## Results

### Improved annotation of *Xcc* strain 8004 genome based on a large transcriptomic dataset

Transcriptomic analyses are intrinsically dependent on the proper structural annotation of genes. In order to improve annotation of *Xcc* strain 8004 using experimental expression data, we produced the transcriptome of two nearly isogenic strains grown in MOKA medium: wild-type strain 8004 and strain 8004::*hrpG*∗ which expresses the constitutive active variant E44K of HrpG (50) (see later for comparative analysis of the HrpG regulon). Total RNAs were extracted from exponentially growing bacteria and subjected to ribodepletion as described (18). Small (<200 nt) and large RNA fractions (>200 nt) were subjected to paired-end and single-end strand-specific sequencing, respectively. RNAs protected by a 5’ triphosphate group in the small fraction were used to map precisely transcriptional start sites. Those 13 libraries corresponding to 88 130 260 and 135 564 422 reads from small and large RNA fractions, respectively, were used to refine the annotation of the genome. More than 1724 transcriptional starts (5’ UTRs) and 1246 3’ UTRs could be experimentally defined (Table 1, https://dx.doi.org/10.25794/reference/id52ofys). Predicted translational start site was modified for 1164 CDS and 753 small ncRNAs were evidenced. These results highlight the importance of experimentally-supported genome annotations and offer improved resources for the functional analysis of *Xcc* transcriptome.

**Table 1.** Impact of transcriptomic data on the *de novo* annotation of *Xcc* strain 8004 genome

| RNaseq data used<sup>a</sup> | All genes | mRNA<sup>b</sup> | rRNA<sup>c</sup> operon | tRNA<sup>d</sup> | Other ncRNA<sup>e</sup> | 5' UTR<sup>f</sup> | 3' UTR<sup>g</sup> | Reference | Accession Number |
| --- | --- | --- | --- | --- | --- | --- | --- | --- | --- |
| No | 4333 | 4273 | 2 | 53 | 1 | - | - | Qian et al., 2005 | NCBI:CP000050.1 |
| No | 4631 | 4478 | 2 | 54 | 93 | - | - | NCBI | NCBI:NC_007086.1 |
| Yes | 5431 | 4617 | 2 | 55 | 753 | 1724 | 1246 | This study | DOI:10.25794/reference/id52ofys |

<sup>a</sup> Use of merged transcriptomic datasets from *Xcc* strain 8004 derivatives (WT or 8004::*hrpG*\*) grown *in vitro* in MOKA medium.
<sup>b</sup> Protein coding sequence
<sup>c</sup> Ribosomal RNA
<sup>d</sup> Transfer RNA
<sup>e</sup> Non coding RNA
<sup>f</sup> 5' untranslated region
<sup>g</sup> 3' untranslated region

### *Xcc* transcriptome remodeling at early stages of hydathode infection

In order to capture a proxy of the physiological status of *Xcc* at the early step of plant leaf infection, we performed RNA sequencing on bacteria re-isolated from cauliflower hydathodes 72 hours after a rapid dip inoculation of an attached leaf. This 72 hpi timepoint corresponds to a biotrophic phase of the infection where bacteria are still limited to the epitemal apoplastic spaces (3). After four hours of continuous immersion of a detached leaf in the bacterial suspension, ca. 10^5^ cfu/hydathode are detected similar to the bacterial titers at 72 hpi (Figure S2). Calculation of Euclidian distances indicated that the 4 hpi transcriptomes cluster with the 72 hpi timepoints rather than *in vitro* samples (Figure S3). This 4 hpi condition was thus chosen as the reference condition since it allows the narrow comparison of two bacterial populations in contact with plant tissues for 4 and 72 hours and a focus on bacterial adaptation to the plant environment.

During hydathode infection (72 hpi versus 4 hpi), *Xcc* massively reshaped its transcriptome with 828 DEGs corresponding to 18% of *Xcc* CDS (Figure 1, Table S4, Table S6). A Gene Ontology (GO) enrichment analysis identified 18 Biological Processes, such as catabolism, stress response and transport, which were significantly enriched in up-regulated genes (Table 2). On the other hand, 13 GO terms were enriched among downregulated genes, with a strong overrepresentation of motility and chemotaxis categories (Table 2).

**Table 2.** GO terms of biological processes enriched among *Xcc* genes differentially expressed in hydathodes (72 hpi vs 4 hpi).

| GO.ID | Term | Annotated<sup>a</sup> | Significant<sup>b</sup> | Expected<sup>c</sup> | Adjust<br>p Value |
| --- | --- | --- | --- | --- | --- |
| **Upregulated** |  |  |  |  |  |
| GO:0006073 | cellular glucan metabolic process | 19 | 11 | 1,89 | 1.8 10<sup>-8</sup> |
| GO:0055114 | oxidation-reduction process | 399 | 74 | 39,78 | 8.9 10<sup>-8</sup> |
| GO:0070814 | hydrogen sulfide biosynthetic process | 5 | 5 | 0,5 | 9.5 10<sup>-6</sup> |
| **GO:0005992** | **trehalose biosynthetic process** | **5** | **5** | **0,5** | **9.5 10<sup>-6</sup>** |
| GO:0006950 | response to stress | 123 | 25 | 12,26 | 0.00063 |
| GO:0019344 | cysteine biosynthetic process | 6 | 4 | 0,6 | 0.00123 |
| GO:0009251 | glucan catabolic process | 11 | 5 | 1,1 | 0.00263 |
| GO:0043649 | dicarboxylic acid catabolic process | 7 | 4 | 0,7 | 0.00265 |
| GO:0044247 | cellular polysaccharide catabolic process | 8 | 4 | 0,8 | 0.00361 |
| GO:0009306 | protein secretion | 46 | 11 | 4,59 | 0.00435 |
| GO:0043623 | cellular protein complex assembly | 18 | 6 | 1,79 | 0.00612 |
| GO:0006457 | protein folding | 18 | 6 | 1,79 | 0.00612 |
| GO:0098661 | inorganic anion transmembrane transport | 15 | 5 | 1,5 | 0.01230 |
| GO:0098869 | cellular oxidant detoxification | 30 | 7 | 2,99 | 0.02468 |
| GO:0044419 | interspecies interaction between organisms | 12 | 4 | 1,2 | 0.02506 |
| GO:0005975 | carbohydrate metabolic process | 201 | 39 | 20,04 | 0.03559 |
| GO:0006817 | phosphate ion transport | 8 | 3 | 0,8 | 0.03753 |
| GO:0016311 | dephosphorylation | 33 | 7 | 3,29 | 0.04006 |
| **Downregulated** |  |  |  |  |  |
| GO:0006935 | chemotaxis | 44 | 36 | 3,12 | < 10<sup>-30</sup> |
| **GO:0097588** | **archaeal or bacterial-type flagellum-dependent cell motility** | **23** | **23** | **1,63** | **9.4 10<sup>-28</sup>** |
| GO:0007165 | signal transduction | 175 | 51 | 12,4 | 1.7 10<sup>-22</sup> |
| **GO:0044781** | **bacterial-type flagellum organization** | **16** | **16** | **1,13** | **2.2 10<sup>-19</sup>** |
| GO:0030031 | cell projection assembly | 16 | 13 | 1,13 | 3.5 10<sup>-13</sup> |
| GO:0000160 | phosphorelay signal transduction system | 133 | 23 | 9,43 | 3.5 10<sup>-5</sup> |
| GO:0000272 | polysaccharide catabolic process | 27 | 7 | 1,91 | 0.0021 |
| GO:0007155 | cell adhesion | 5 | 3 | 0,35 | 0.0031 |
| **GO:0048870** | **cell motility** | **25** | **25** | **1,77** | **0.0039** |
| GO:0071554 | cell wall organization or biogenesis | 40 | 6 | 2,83 | 0.0149 |
| GO:0006468 | protein phosphorylation | 79 | 11 | 5,6 | 0.0215 |
| GO:0015031 | protein transport | 81 | 12 | 5,74 | 0.0295 |
| GO:0000302 | response to reactive oxygen species | 11 | 3 | 0,78 | 0.0378 |

<sup>a</sup> Number of genes annotated for a given GO term
<sup>b</sup> Number of differentially regulated genes among the genes annotated for a given GO term
<sup>c</sup> Expected number of differentially regulated genes if no significant enrichment
<sup>e</sup> Terms for which all annotated genes are differentially regulated are highlighted in bold

**Figure 1.**
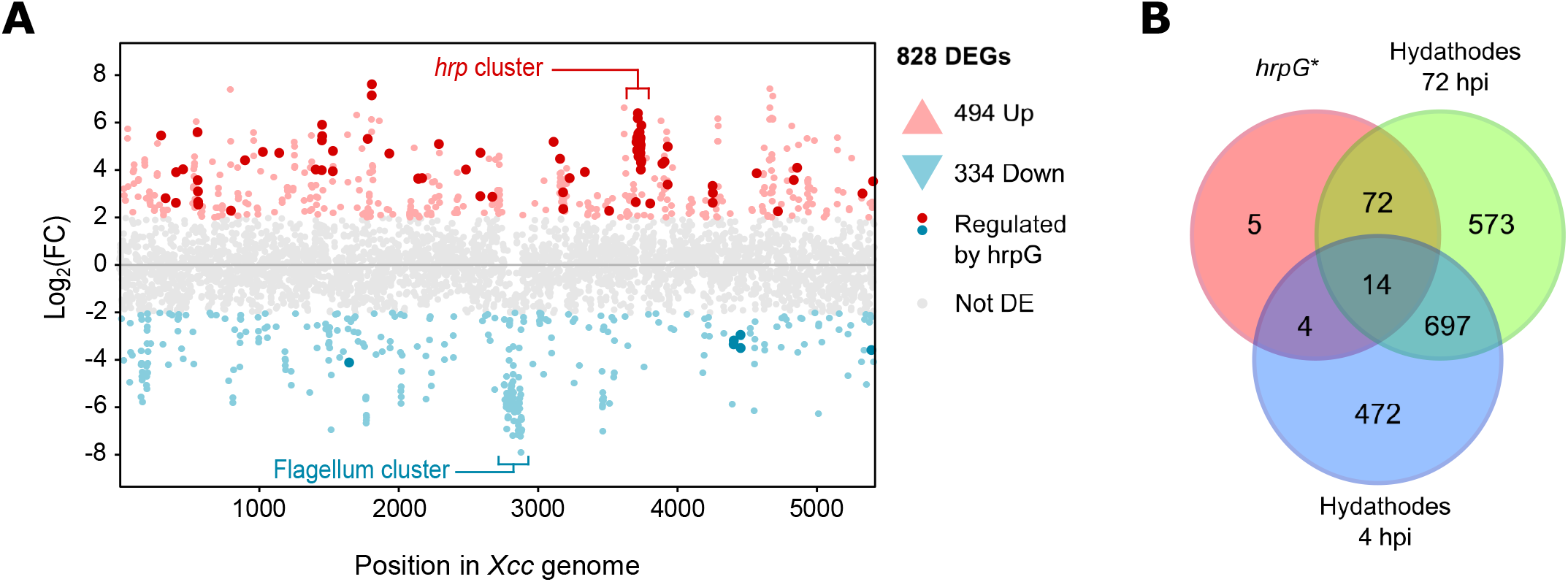
*Xanthomonas campestris* undergoes a massive transcriptomic reprogramming during hydathodes infection. (A) Genome-wide expression profile of *Xcc* in hydathodes at 72 hpi versus 4 hpi. Each point represents a gene for which the change in expression level is given as the log_2_ fold change (Log_2_FC) between 72 hpi and 4 hpi into hydathodes. Genes considered significantly differentially expressed (DEGs) are represented in red if induced or in blue if repressed between the two timepoints. Non DEGs are colored in grey. Genes under the control of the HrpG regulator (*ie*. found differentially expressed in the 8004::*hrpG*∗ vs 8004 dataset) and found differentially expressed at 72 hpi vs 4 hpi in hydathodes are colored in dark red (up-(A) regulated) and dark blue (down-regulated). (B) Venn diagram showing the total number of DEGs (|Log_2_FC| ≥ 2; FDR ≤ 0.05) obtained after growth of the *Xcc* 8004 WT strain in MOKA-rich medium as compared to either the 8004::*hrpG*∗ mutant in MOKA, the WT strain after 4 hours into hydathodes or 72 hours into hydathodes.

### *Xcc* adopts a sedentary lifestyle inside hydathodes

Expression of most genes coding for biosynthesis of flagella and type IV pili and chemotaxis are strongly repressed at 72 hpi. Motility and chemotaxis are key components of pathogenicity, especially for plant pathogens (51). However, *Xcc* seems not to be flagellated when growing in xylem fluids and the motile *Xcc* cells seemed less pathogenic on cauliflower and radish (52). These observations suggest a probable fitness cost of motility during infection. To investigate the importance of motility in disease development, we constructed mutants in key genes for the synthesis of type IV pilus (Δ*pilE* and Δ*pilA*) or flagella (Δ*fliC* and Δ*fliQ*). Single and multiple mutants were tested for *in vitro* motility (Figure S4), pathogenicity (Figure S5) and hydathode colonization (Figure 2). Despite expected *in vitro* motility phenotypes (Figure S4), none of the tested mutant affected disease symptoms development nor hydathode colonization (Figure S5, Figure 2) as it might have been expected for genes whose expression is repressed inside hydathodes. Altogether, these results suggest that motility is a process that is not needed for hydathode infection and maybe costly at this stage of the infection.

**Figure 2.**
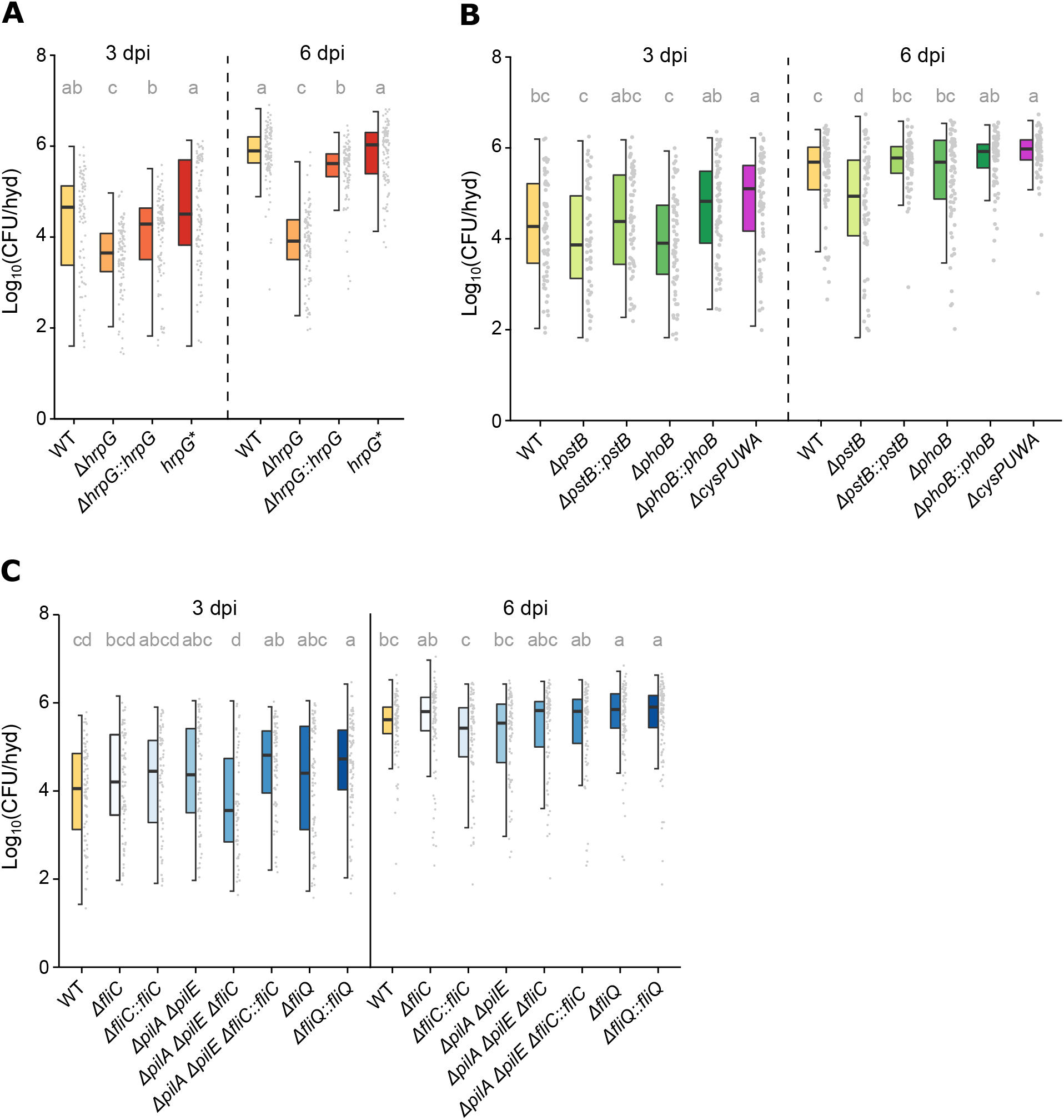
*Xanthomonas* colonization of hydathodes. Bacterial multiplication of *Xcc* 8004 wild-type strain (WT), deletion mutants and complemented strains in individual hydathodes 3 and 6 days after dip-inoculation of the second true leaf of 4 weeks-old cauliflower plants. The box plot representations are showing the impact of (A) mutations in the T3SS *hrpG* regulator, (B) mutations in phosphate and sulfate transport genes and, (C) mutations in motility genes over *Xcc* multiplication into hydathodes. Each point of the plot represents the population extracted from one hydathode. At least eight </i> hydathodes were sampled on one leaf per plant and three plants were used per experiment, though not all hydathodes were infected. Results from at least three independent experiments were pooled and a total of at least 50 infected hydathodes were counted for each strain. Letters indicate statistically different groups obtained from the Kruskal-Wallis test on all data points for each strain with an error α = 0.05.

### Activation of the HrpG regulon at early steps of hydathode infection

*hrpG* gene is a known master regulator required for the expression of the T3S machinery, T3E proteins and additional genes including plant cell wall degrading enzymes (PCWDE) in *Xanthomonas* spp.. In *Xcc*, expression of *hrpG* and 49 (out of 55) genes coding for the T3S system and type 3-secreted proteins, was induced at 72 hpi suggesting the involvement of HrpG at this stage of infection (Table S4 and S7). We thus investigated the biological importance of *hrpG* during hydathode infection and studied its regulon.

The 8004Δ*hrpG* mutant showed a 10-fold reduced multiplication in hydathodes at 72 hpi (Figure 2A) and was avirulent on cauliflower after wound inoculation (Figure S5). Both phenotypes could be complemented (Figure 2A and S5). The *hrpG*∗ (E44K) gain-of-function mutation conferring constitutive expression of the HrpG regulon *in vitro* did not affect the multiplication of *Xcc* in cauliflower hydathodes (Figure 2A) nor its pathogenicity after wound inoculation relative to the wild-type strain. As observed in *Xanthomonas euvesicatoria* (20), the 8004::*hrpG*∗ strain also presented a reduced extracellular protease activity (Figure S6B). This suggests that HrpG∗ *in planta* functions are retained and that *in vitro* studies with this mutant are legitimate.

To determine the extent of the HrpG regulon, we compared the transcriptomes of the 8004::*hrpG*∗ and wild-type strains grown in MOKA medium (Table S4, Table S6). Wild-type strain 8004 does not express *hrp* genes in MOKA in contrast to strain 8004::*hrpG*∗. Analysis of the HrpG regulon identified 95 DEGs (Log2(fold change) ≥ 2 or ≤ −2, FDR adjusted *p*-value < 0.05) (Table S4). Among the 85 genes with an increased expression in strain 8004::*hrpG*∗, 43 possess a PIP box promoter motif and are thus likely under *hrpX* control. 39 of the 95 DEGs correspond to genes involved in T3SS and T3Es explaining the enrichment in the GO term “secretion” among genes upregulated in strain 8004::*hrpG*∗ (Table 3, Table S4 and S7). Expression of only 18 out of the 30 genes encoding type 3 secreted proteins was increased in strain 8004::*hrpG*∗ compared to 24 genes at 72 hpi inside hydathodes. Among the other 56 DEGs of the HrpG regulon, 26 encode proteins with unknown function and 19 PCWDE.

**Table 3.**
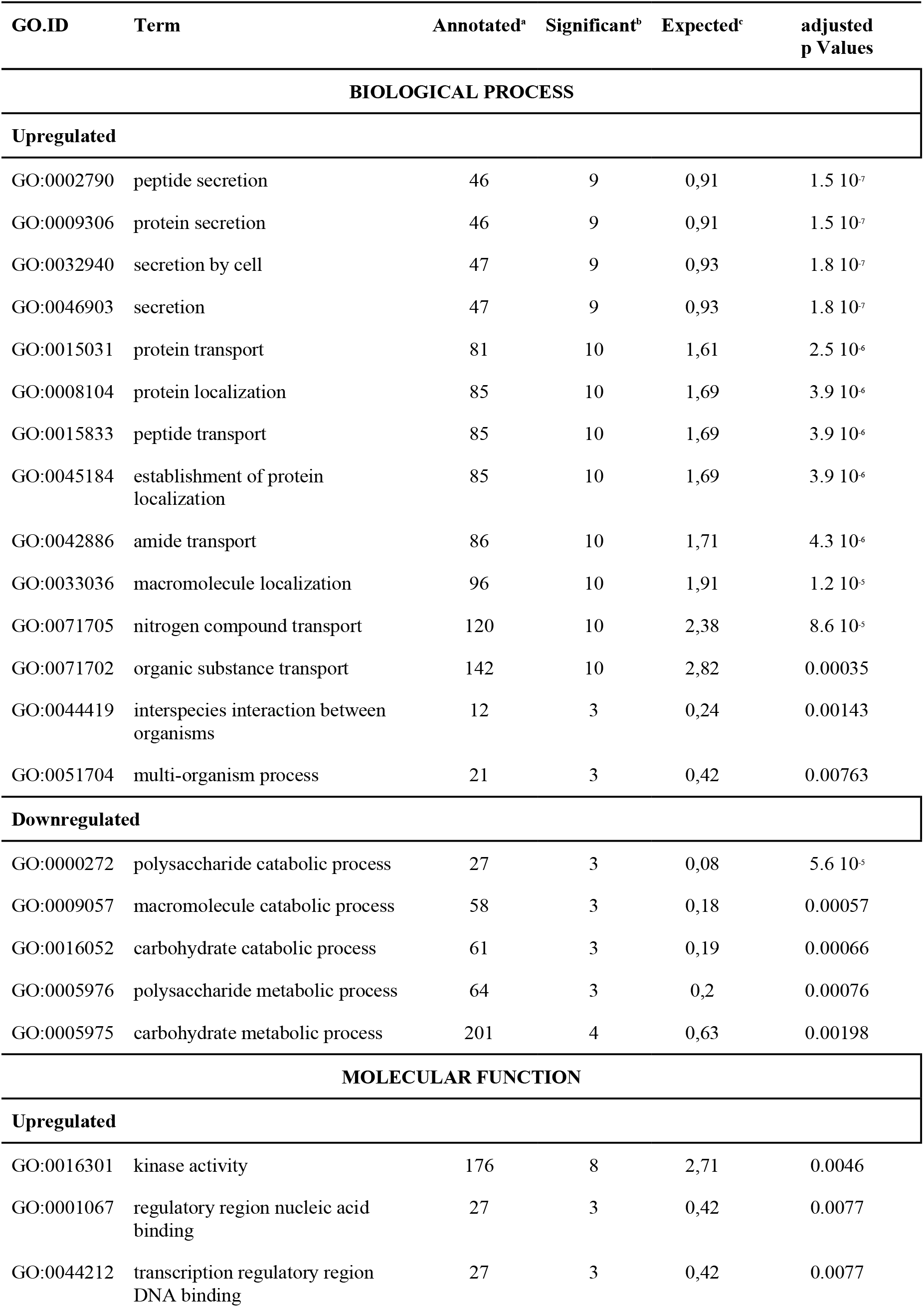
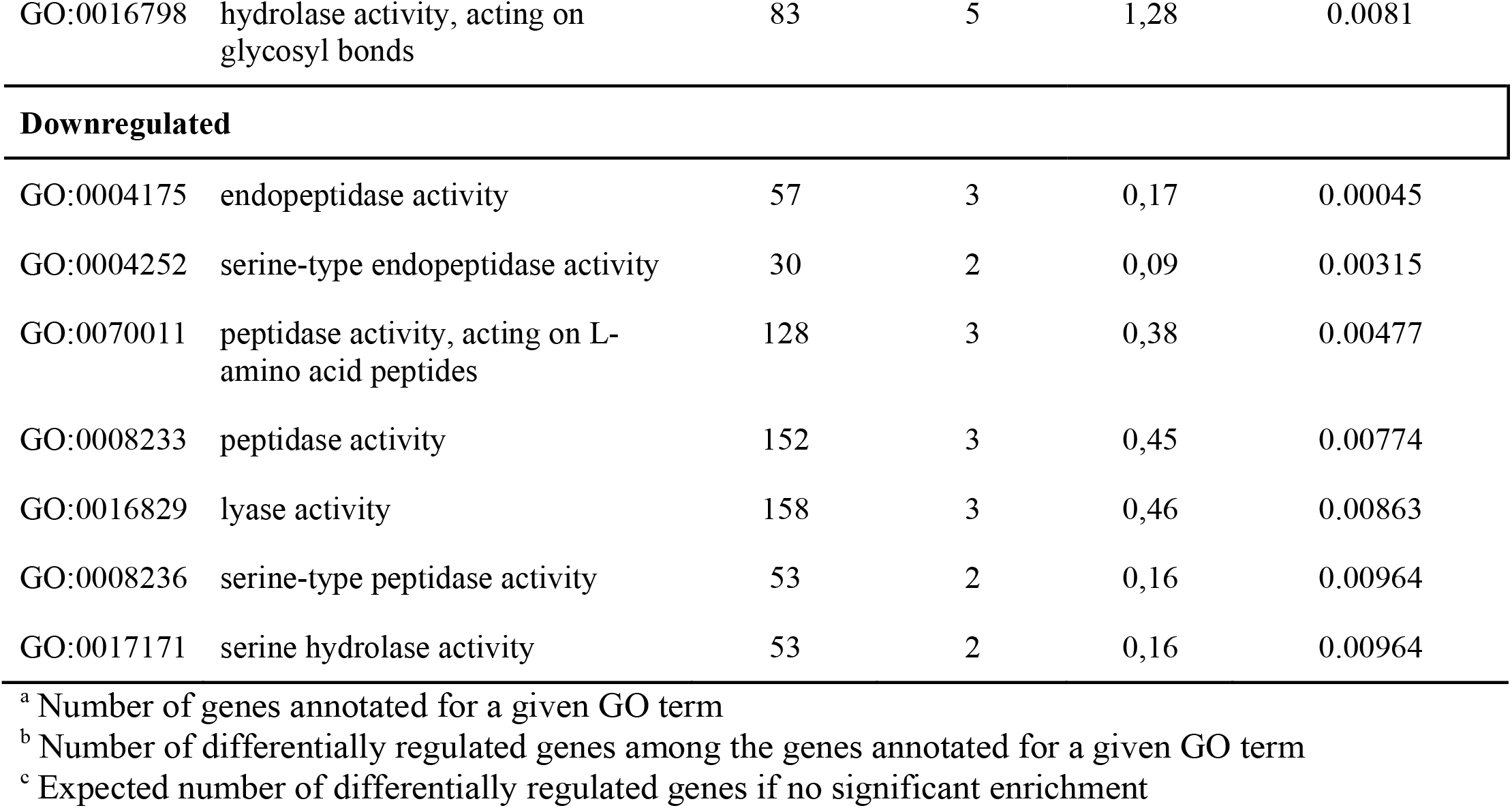
GO terms of biological processes enriched among *Xcc* genes belonging to the HrpG regulon.

| GO.ID | Term | Annotated<sup>a</sup> | Significant<sup>b</sup> | Expected<sup>c</sup> | adjusted p Values |
| --- | --- | --- | --- | --- | --- |
| **BIOLOGICAL PROCESS** |  |  |  |  |  |
| **Upregulated** |  |  |  |  |  |
| GO:0002790 | peptide secretion | 46 | 9 | 0,91 | 1.5 10<sup>-7</sup> |
| GO:0009306 | protein secretion | 46 | 9 | 0,91 | 1.5 10<sup>-7</sup> |
| GO:0032940 | secretion by cell | 47 | 9 | 0,93 | 1.8 10<sup>-7</sup> |
| GO:0046903 | secretion | 47 | 9 | 0,93 | 1.8 10<sup>-7</sup> |
| GO:0015031 | protein transport | 81 | 10 | 1,61 | 2.5 10<sup>-6</sup> |
| GO:0008104 | protein localization | 85 | 10 | 1,69 | 3.9 10<sup>-6</sup> |
| GO:0015833 | peptide transport | 85 | 10 | 1,69 | 3.9 10<sup>-6</sup> |
| GO:0045184 | establishment of protein localization | 85 | 10 | 1,69 | 3.9 10<sup>-6</sup> |
| GO:0042886 | amide transport | 86 | 10 | 1,71 | 4.3 10<sup>-6</sup> |
| GO:0033036 | macromolecule localization | 96 | 10 | 1,91 | 1.2 10<sup>-5</sup> |
| GO:0071705 | nitrogen compound transport | 120 | 10 | 2,38 | 8.6 10<sup>-5</sup> |
| GO:0071702 | organic substance transport | 142 | 10 | 2,82 | 0.00035 |
| GO:0044419 | interspecies interaction between organisms | 12 | 3 | 0,24 | 0.00143 |
| GO:0051704 | multi-organism process | 21 | 3 | 0,42 | 0.00763 |
| **Downregulated** |  |  |  |  |  |
| GO:0000272 | polysaccharide catabolic process | 27 | 3 | 0,08 | 5.6 10<sup>-5</sup> |
| GO:0009057 | macromolecule catabolic process | 58 | 3 | 0,18 | 0.00057 |
| GO:0016052 | carbohydrate catabolic process | 61 | 3 | 0,19 | 0.00066 |
| GO:0005976 | polysaccharide metabolic process | 64 | 3 | 0,2 | 0.00076 |
| GO:0005975 | carbohydrate metabolic process | 201 | 4 | 0,63 | 0.00198 |
| **MOLECULAR FUNCTION** |  |  |  |  |  |
| **Upregulated** |  |  |  |  |  |
| GO:0016301 | kinase activity | 176 | 8 | 2,71 | 0.0046 |
| GO:0001067 | regulatory region nucleic acid binding | 27 | 3 | 0,42 | 0.0077 |
| GO:0044212 | transcription regulatory region DNA binding | 27 | 3 | 0,42 | 0.0077 |
| GO:0016798 | hydrolase activity, acting on glycosyl bonds | 83 | 5 | 1,28 | 0.0081 |
| **Downregulated** |  |  |  |  |  |
| GO:0004175 | endopeptidase activity | 57 | 3 | 0,17 | 0.00045 |
| GO:0004252 | serine-type endopeptidase activity | 30 | 2 | 0,09 | 0.00315 |
| GO:0070011 | peptidase activity, acting on L-amino acid peptides | 128 | 3 | 0,38 | 0.00477 |
| GO:0008233 | peptidase activity | 152 | 3 | 0,45 | 0.00774 |
| GO:0016829 | lyase activity | 158 | 3 | 0,46 | 0.00863 |
| GO:0008236 | serine-type peptidase activity | 53 | 2 | 0,16 | 0.00964 |
| GO:0017171 | serine hydrolase activity | 53 | 2 | 0,16 | 0.00964 |

<sup>a</sup> Number of genes annotated for a given GO term
<sup>b</sup> Number of differentially regulated genes among the genes annotated for a given GO term
<sup>c</sup> Expected number of differentially regulated genes if no significant enrichment

The HrpG regulon was almost entirely included in the *in planta* transcriptome (90 out of 95 genes) indicating that HrpG is activated during hydathode infection (Figure 1). Eighteen genes of the HrpG regulon are overexpressed at 4 hpi in comparison to the MOKA condition indicating that expression of the T3S machinery is initiated early during infection (Table S5). Yet, the HrpG regulon (95 genes) remains a very minute part of *Xcc* transcriptomic adaptations to “*in planta”* conditions (828 genes).

### Metabolic adaptations of *Xcc* inside hydathodes highlight several nutritional properties of the plant environment

Among the 31 GO terms significantly affected at transcriptomic level, a third are involved in metabolism, indicating that *Xcc* undergoes an important metabolic adaptation inside the hydathode.

#### Modification of the expression profile of genes encoding PCWDE

We observe an increased transcription of genes involved in the catabolism of cellulose, a major component of primary cell wall. Yet, increased expression of other PCWDE genes is not observed at 72 hpi, consistently with the absence of visible degradation of cell walls in the epithem (3). Among the 43 genes encoding PCWDE in *Xcc* strain 8004 (53), 10 have an increased and 10 a reduced expression at 72 hpi, respectively (Table S7-2). Expression of either *xps* or *xcs* genes encoding type II secretion systems involved in PCWDE secretion is not induced at 72 hpi suggestive of a biotrophic lifestyle.

Lignin is another major component of plant cell walls and a source of aromatic compounds. Interestingly, the expression of *XC_3426* and X*C_3427* genes coding for protocatechuate (PCA) 4;5-dioxygenase subunits is increased 6 and 8 fold at 72 hpi, respectively. PCA is a lignin degradation product (54) which can be further catabolized by PCA dioxygenases to enter the tricarboxylic acid (TCA) cycle (55). While *XC_3426* and *XC_3427* relevance for pathogenicity remains unknown, *XC_0375* to *XC_0383* genes cluster encoding 3-and 4-hydroxybenzoate degrading enzymes are needed for full virulence of *Xcc* in radish (55) suggesting that degradation of plant phenolic compounds happens inside plant tissues.

#### Expression of transporter genes is deeply remodeled *in planta*

Broad expression changes can be observed in transporter genes since 49 out of 210 genes involved in transport are differentially expressed at 72 hpi (Table 2, Table S7-1). Those transporters belong to MFS, ABC and TBDT families. TBDTs have been shown to be involved in iron and carbohydrates polymers uptake with high affinity in *Xcc* (14, 38, 56). 14 out of 48 TBDT genes are differentially expressed: 7 show an increased expression at 72 hpi and 7 with a decreased activity. While most have no known function, those which expression is induced by polygalacturonate (PGA) are less expressed at 72 hpi (Table S7-1) (14). The two TBDT genes *XC_3205* and *XC_2512* known to be positively regulated by HrpG and HrpX (14) are induced at 72 hpi.

Interestingly, the absence of the *fur* regulon and its iron high affinity TBDT transporters (*XC_0167, XC_3463, XC_2846, XC_1341, XC_1108 XC_0924-0925, XC_0642, XC_4249, XC_0558* and *XC_4249*) in our dataset indicates that iron might not be limiting at this stage of hydathode infection.

#### Upregulation of two pathways important for sulfur assimilation in hydathodes

In contrast to iron, induction of genes important for sulfur transport and assimilation is observed at 72 hpi in hydathodes: the operon encoding the ABC sulfate transporter CysPUWA (*XC_3292* to *XC_3295*) and the operon encoding an assimilatory sulfate reduction pathway leading to sulfide production (*XC_0990* to *XC_0994*) are induced by 6 and 20 fold, respectively (Table S4). Sulfide is then available for cysteine and methionine biosynthesis. Sulfur metabolism including sulfur-containing amino acids, sulfur compounds, or sulfate have been shown to be involved in different virulence factor production, as in xanthan production (57) or T3SS induction (58). Yet, a deletion of the entire *cysPUWA* operon (Δ*XC_3292-95*) in *Xcc* strain 8004 did not significantly affect bacterial multiplication neither in hydathodes after dip-inoculation nor disease symptom development after wound inoculation in cauliflower (Figure 2, Figure S5). Thus, sulfate import through the CysPUWA system is not limiting for bacterial growth in hydathodes or sulfur might be acquired through other import pathways such as the taurine import system. *E. coli* responds to sulfate or cysteine starvation by expressing the *ssuABCDE* and *tauABCD* operons which are involved in the uptake of alkanesulfonate and desulfonation of the organosulfonates and for uptake and desulfonation of taurine, respectively (59, 60). While *ssuABCDE* is absent in *Xcc* strain 8004, TauABCD homologues are encoded by genes of the locus *XC_3454* to *XC_3460* (Figure S7). Interestingly, expression of these genes is increased at 72 hpi suggesting a possible implication of this pathway in sulfur assimilation *in planta*.

#### Phosphate uptake machinery is rate limiting for *Xcc* multiplication inside hydathodes

Increased expression of the genes encoding the PstSCAB high-affinity transporter system (genes *XC_2708* to *XC_2711*, Table S4) involved in active inorganic phosphate (Pi) import upon phosphate starvation suggests that Pi might be limiting inside hydathodes. This system is known to be activated in various conditions in bacteria (61), including during plant colonization, and is essential for *Xanthomonas axonopodis* pv. *citri* (*Xac*) pathogenicity on citrus (62, 63). In order to test if phosphate acquisition is important for *Xcc* strain 8004 during hydathode colonization, we mutated *XC_2711* (*pstB*) and *XC_3272* which encodes an homologue of the PhoB response regulator important for *E. coli* Pi starvation response (64, 65). Results obtained demonstrated that *pstB*, unlike *phoB*, is important for hydathode colonization (Figure 2B) and that both Δ*pstB* and Δ*phoB* mutants caused symptoms similar to the wilt-type strain after direct inoculation into xylem vessels (Figure S5B). These results demonstrate that Pi might be limiting specifically for the growth in hydathodes. Similar to Pi, expression of genes important for nitrogen assimilation such as those involved in uptake of nitrite and nitrate and their reduction to ammonia (*XC_2175* to *XC_2178*) are induced at 72 hpi, further stressing the importance of *Xcc* mineral nutritional needs in hydathodes.

### *Xcc* adapts to nutritional, osmotic and environmental stresses in hydathodes

Surprisingly, there are limited transcriptional changes for genes associated with transcription, translation, replication, TCA cycle or amino acid biosynthesis between 4 and 72 hpi. However, these functions are already strongly repressed at 4 hpi compared to MOKA conditions indicating that adaptation to the plant environment is associated with a rapid repression of cellular division and core-metabolism. Other stress-responsive genes have an increased expression *in planta* such as base excision and nucleotide excision repair systems, trehalose production pathways, superoxide dismutases and chaperone proteins. In addition, expression of the ABC transporter system *OpuB-ABC* (66, XC_0173 to XC_0174) and the choline degradation pathway (67, XC_0760 to XC_0761), both involved in osmoprotection, are induced 72 hpi suggesting that bacteria face an osmotic shock in the apoplast of the epithem cells (Figure 3, Table S4). Finally, expression of some genes of the *gum* operon (*XC_1658* to *XC_1673*) implicated in the production of the xanthan exopolysaccharides (EPS) are induced by 4-to 5-fold at 72 hpi. Xanthan is a well-known protectant against environmental stresses and toxic compounds and a suppressor of plant immunity (68).

**Figure 3.**
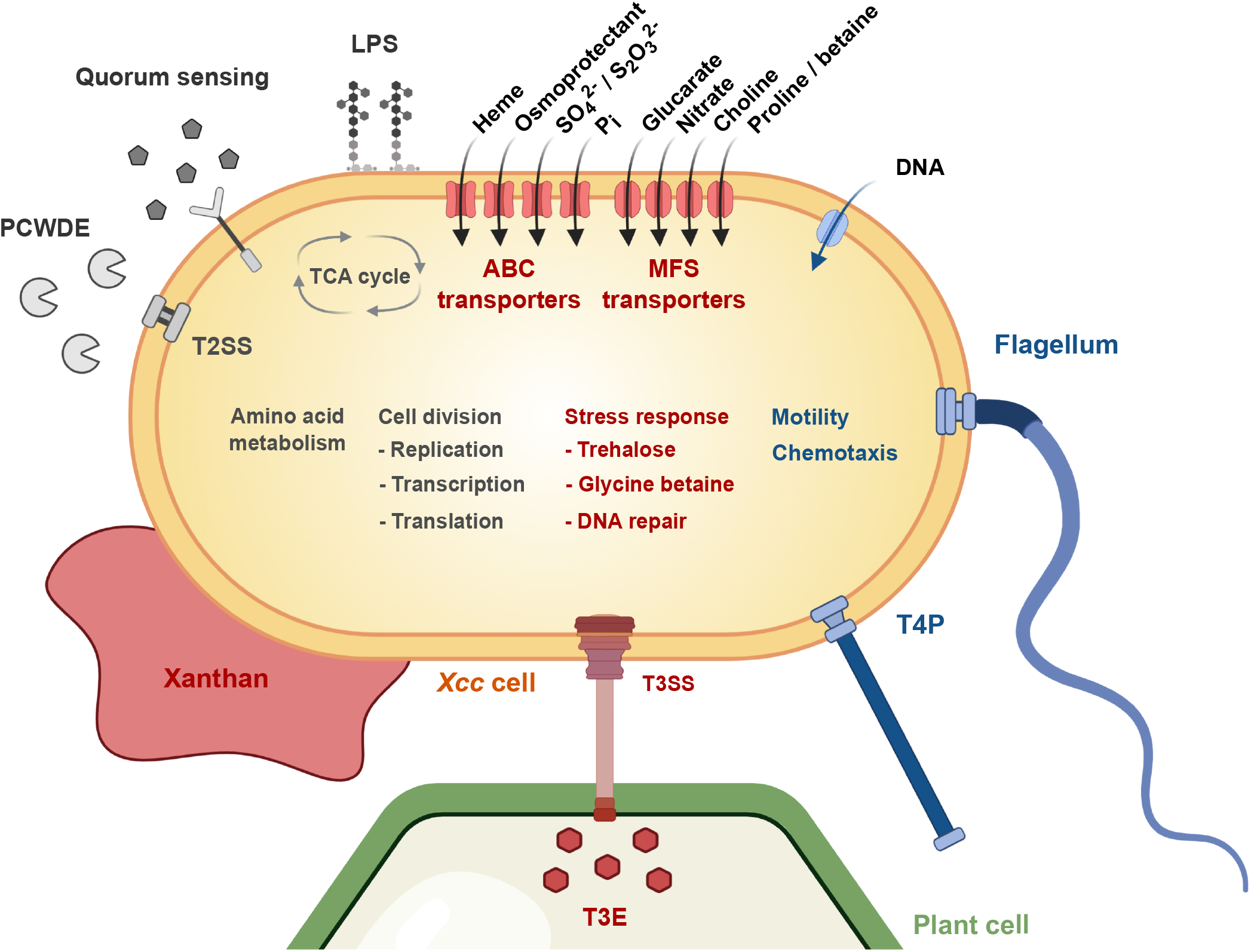
Schematic representation of *Xcc* main transcriptomic responses happening during the early step of hydathode infection. Genes corresponding to blue and red objects are repressed and induced between 4 and 72 hpi, respectively. Genes corresponding to grey objects are not differentially expressed. T3SS: type three secretion system; T3E: type three effector; T2SS: type two secretion system; LPS: lipopolysaccharide; PCWDE: Plant cell wall degrading enzymes; T4P: type four pilus. Figure drafted using biorender (https://app.biorender.com).

Altogether, these data seem to indicate that *Xcc* cells must cope with some nutritional, osmotic and environmental stresses in hydathodes.

## Discussion

*Xcc* life cycle depends on its adaptation to various plant environments (e.g. seeds, leaf surface, hydathodes, xylem, mesophyll, debris). This work describes the transcriptional changes that *Xcc* undergoes upon plant infection and hydathode colonization, including the regulation of various metabolic and virulence pathways (Figure 3) and informs about the environmental conditions faced by *Xcc* inside hydathodes.

### *Xcc* adopts a sedentary biotrophic lifestyle inside hydathodes

*Xcc* is a known necrotroph causing black rot disease. However, the physiological snapshot obtained by RNAseq at 72 hpi in hydathodes suggests that *Xcc* behaves as a biotroph since the epithem is intact (69) and many genes coding for degradative enzymes, T2SS and most catabolic pathways of sugar polymers and carbohydrates were expressed at low levels. We also observed that the constitutive activation of the HrpG regulon recapitulating part of the *in planta* condition at 72 hpi is associated with a reduced extracellular protease activity. *Xcc* carbon and nitrogen needs could be supported *in planta* by the continuous flow of xylem sap in the epithem. Furthermore, the expression of multiple virulence-associated genes such as those involved in quorum sensing, iron uptake or motility was also reduced at 72 hpi in hydathodes. Though motility has been shown to be an essential virulence trait for many bacterial pathogens (70), *Xcc* motility mutants were not affected in pathogenicity as expected from genes whose expression is repressed in hydathodes. Repression of chemotaxis and motility has also been reported at early rice infection stages in *Xanthomonas oryzae pv. oryzicola* (*Xoc*) (34) while twitching motility and quorum sensing were both activated at later infection stages and important for virulence of *Xanthomonas oryzae pv. oryzae* (*Xoo*) (32) and *Xac* (28). Interestingly, an increase in the expression of genes important for twitching motility and adhesion was observed in *Xcc* grown *in vitro* in xylem sap (13). Xylem sap corresponds to the environment met by *Xcc* immediately after leaving the hydathode. These observations suggest a possible biphasic infectious process with the reactivation of the motility and other virulence-associated genes at later infection stages. Further transcriptomic analyses of virulence gene expression throughout the entire infectious cycles would be needed to support such a hemibiotrophic lifecycle of *Xcc*.

### Stealthiness of *Xcc* inside hydathodes and neutralization of plant immune responses

The observed repression of chemotaxis, twitching and swimming motility also suggests that those functions are dispensable if not detrimental once inside hydathodes. For instance, production of bacterial peptide flg22 from the flagellar FliC protein is a well-known PAMP (pathogen-associated molecular pattern) recognized by the FLS2 receptor and a potent elicitor of basal plant immunity (71). While flg22_8004_ peptide is not recognized by Arabidopsis FLS2 (72, 73), we cannot exclude that other FliC peptides, flagellar proteins or pili proteins from *Xcc* strain 8004 could act as PAMPs in Brassicaceae. In the absence of an *Xcc* strain constitutively expressing those genes, we were not able to test such hypotheses. *Xcc* stealthiness could also be acquired by limiting bacterial multiplication until an efficient suppression of immunity has been achieved. In contrast to a wild-type *Xcc* strain, a T3S system mutant unable to deliver T3E proteins inside plant cells caused hydathode browning and necrosis at 48 hpi and had a reduced multiplication at 72 hpi (3). These results indicate that hydathode immune responses can be effective against bacterial pathogens and that their suppression by *Xcc* T3S system and its T3E proteins is required for successful infection.

### Inference of environmental conditions inside hydathodes based on *Xcc* transcriptomic behaviour

Compared to *in vitro*-grown *Xcc*, transcriptomic changes are already observed as early as 4 hpi with the increased expression of 14 genes belonging to HrpG regulon (Table S4, Table S5). These genes could participate in the transition from *in vitro* to *in planta* growth such as gene XC_2566 coding for an extracellular function (ECF) sigma factor which importance for this transcriptional switch could be tested. Those transcriptomic profiles can also be used to infer the metabolic and physiological status of *Xcc* inside hydathodes and the nutritional properties of the epithem. For instance, the epithem environment is likely not limiting for assimilable iron since the corresponding uptake machinery is not expressed. In contrast, both the low-affinity phosphate inorganic transport (Pit) and the high-affinity phosphate-specific transport (Pst) systems important for the uptake of inorganic phosphate (Pi) in *Xanthomonas* are upregulated at 72 hpi: (62, 74). Similar to *Xac* (62), *Xcc* Pst system is needed for growth *in planta*. These results demonstrate that *Xcc* not only faces Pi starvation inside hydathodes but that Pi availability also limits *Xcc* proliferation in this tissue. Very low Pi concentrations are indeed found in guttation fluids of several plant species such as barley (75) and are correlated with expression of genes coding for plant high-affinity Pi transporters such as *AtPHT1;4* in hydathodes even under Pi-sufficient conditions (76). These observations suggest that an active competition between the plant and *Xcc* for access to Pi occurs in the epithem. Similar to Pi, *Xcc* transcriptome at 72 hpi also suggests that sulfur and nitrogen are present in low amounts requiring the upregulation of dedicated uptake systems. Yet, it remains unclear whether these elements are limiting for growth of *Xcc* inside hydathodes. Exposure to stresses is also unveiled by the transcriptomic upregulation of genes involved in responses to general stress (e.g. *gum* genes) and osmotic stress. However, it remains uncertain whether osmotic stress is intrinsic of the epithemal environment or whether it is caused by plant immunity. For instance, we could not evidence significant signs of oxidative stresses classically associated with strong plant immune responses. Therefore, *Xcc* seems to adapt rapidly to the low concentrations of nutrients found in the epithem and to endure limited stress maybe due to the continuous flow of fluids inside hydathodes which renews nutrient supplies and dilutes potential antibacterial compounds.

Such global transcriptomic study provides an averaged picture of the bacterial population *in planta* and will feed functional genomic approaches of *Xcc* pathogenicity.

## Acknowledgments

AC and JL were funded by a PhD grant from the French Ministry of Higher Education, Research and Innovation. BR, AC and LDN were funded by grants from the Agence Nationale de la Recherche XANTHOMIX (ANR-2010-GENM-013-02), Xopaque (ANR-10-JCJC-1703-01) and NEPHRON (ANR-18-CE20-0020-01). We are grateful to Stéphanie Bolot for management and early analysis of the Xanthomix dataset. JL, EL, AB and LDN were funded by the XBOX (ANR-19-CE20-JCJC-0014-01) project. The Laboratoire des Interactions Plantes-Microorganismes is part of the French Laboratory of Excellence project (TULIP ANR-10-LABX-41; ANR-11-IDEX-0002-02). All authors benefited from the COST action CA16107 EuroXanth.

## Conflict of interest

The authors declare no conflict of interest.

## Supplemental Tables

**Table S1:** List of strains and vectors used in this study

**Table S2:** Sequences of oligonucleotides used for construction of deletion and complementation plasmids

**Table S3:** Sequences of oligonucleotides used for oligocapture of plant RNAs

**Table S4:** Differentialy expressed genes in each condition tested.

**Table S5:** List of genes belonging to HrpG regulon already differentially expressed after 4 hpi into hydathodes.

**Table S6:** Complete results of the RNAseq experiments.

**Table S7:** Summary of *in planta* RNAseq results sorted for specific biological functions.

**Table S8:** Properties of RNAseq libraries from *Xcc* strain 8004 wild-type and derivatives grown *in vitro* or harvested from cauliflower hydathodes.

## Supplemental Figures

**Figure S1:**
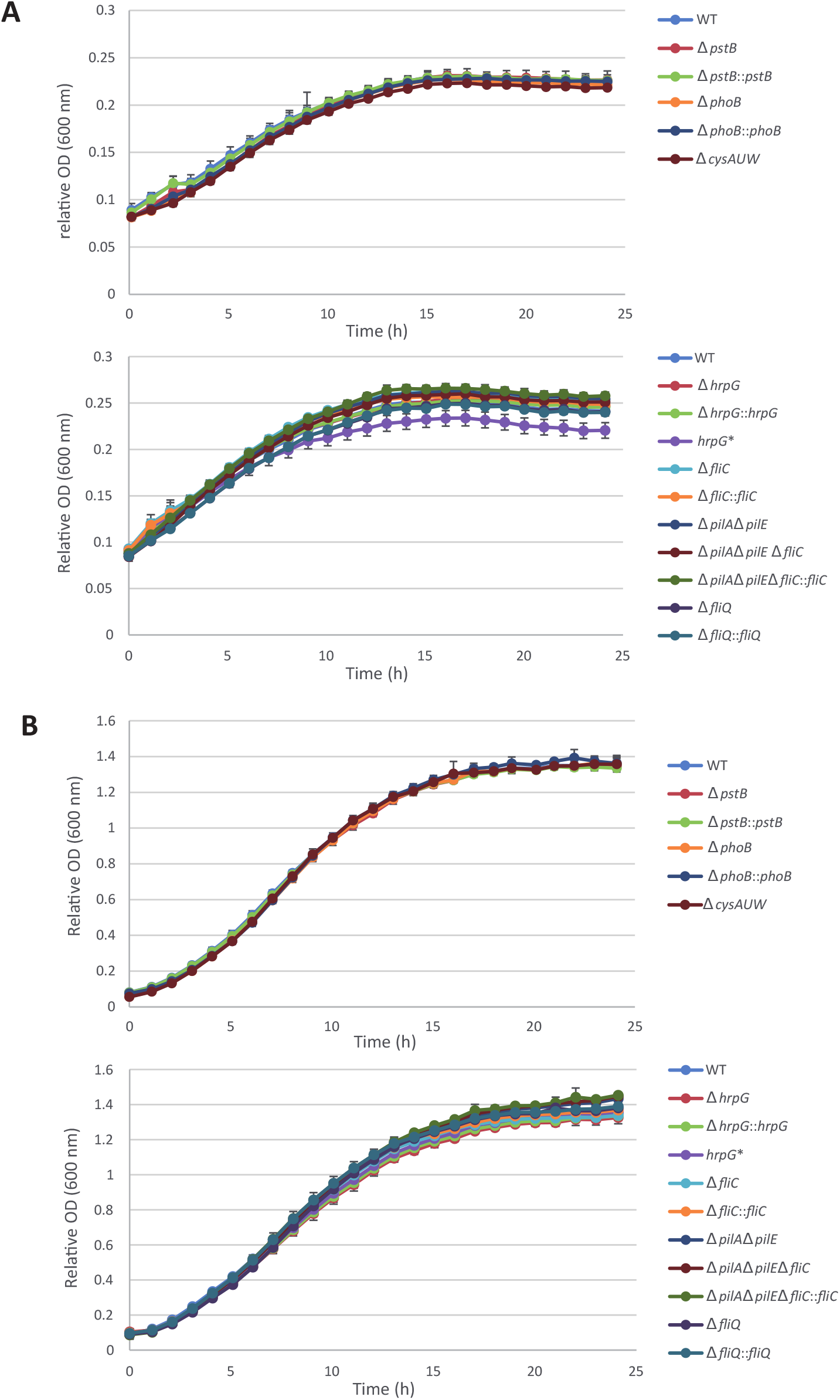
Growth of Xcc wild-type and mutant strains in MME minimal medium (A) and MOKA rich (B) medium. After overnight growth in complete medium, cells were harvested, washed, and resuspended in MME or MOKA. The error bars indicate the standard deviations obtained from 4 technical replicates. The experiments were repeated 3 times an d similar results were obtained.

**Figure S2.**
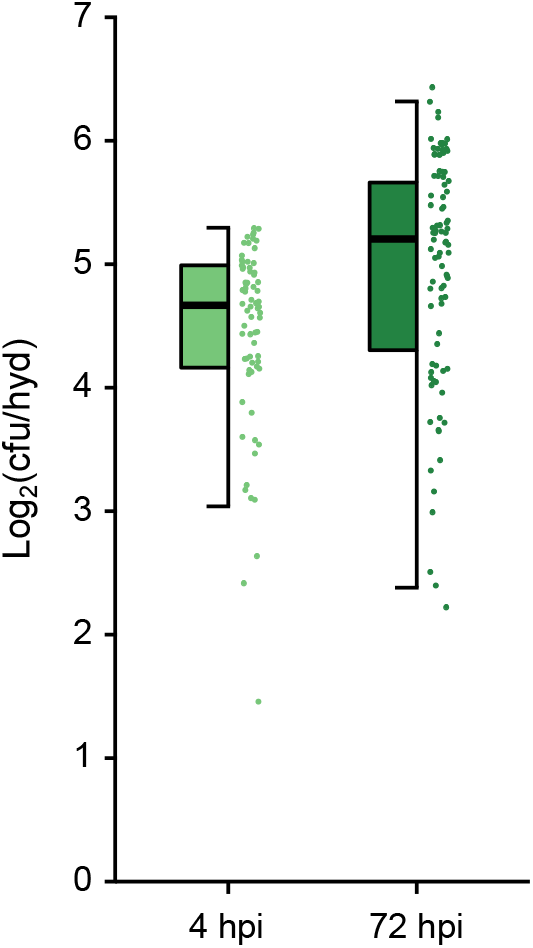
Measure of hydathode population in the two conditions used for RNAseq. The number of *Xcc* colony forming units (cfu) per hydathode (hyd) was determined after 4 hours of continuous dipping (4 hpi) or 72 hours after transient dipping (72 hpi) for 24 or 30 individual hydathodes respectively in three independent biological replicates. Each point of the plot represents the population extracted from one hydathode. Box plots represent the pooled results obtained from three independent experiments.

**Figure S3.**
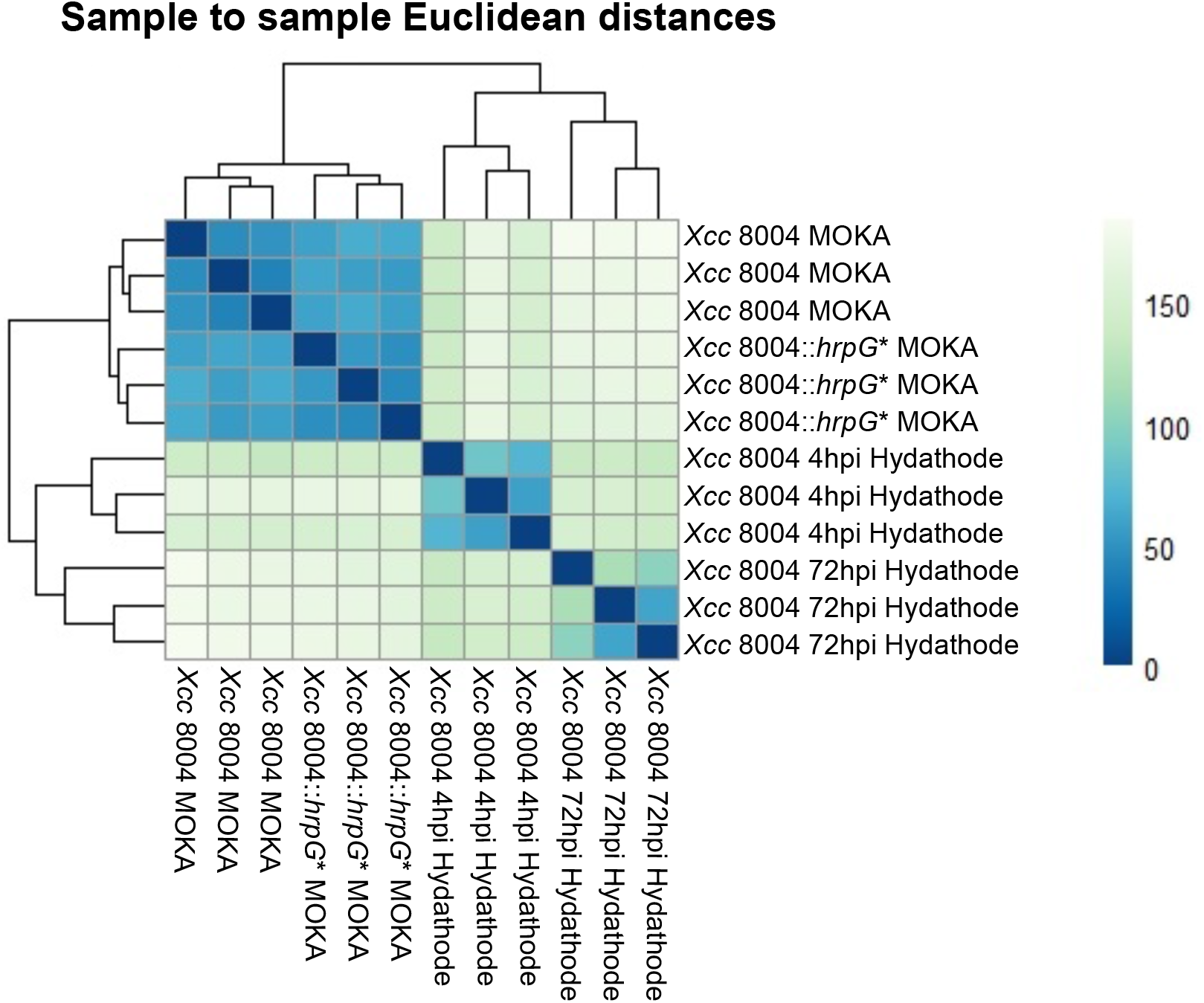
RNAseq samples clustering. The heatmap shows the Euclidian distances between samples as calculated from the variance-stabilizing data transformation of the count data.

**Figure S4.**
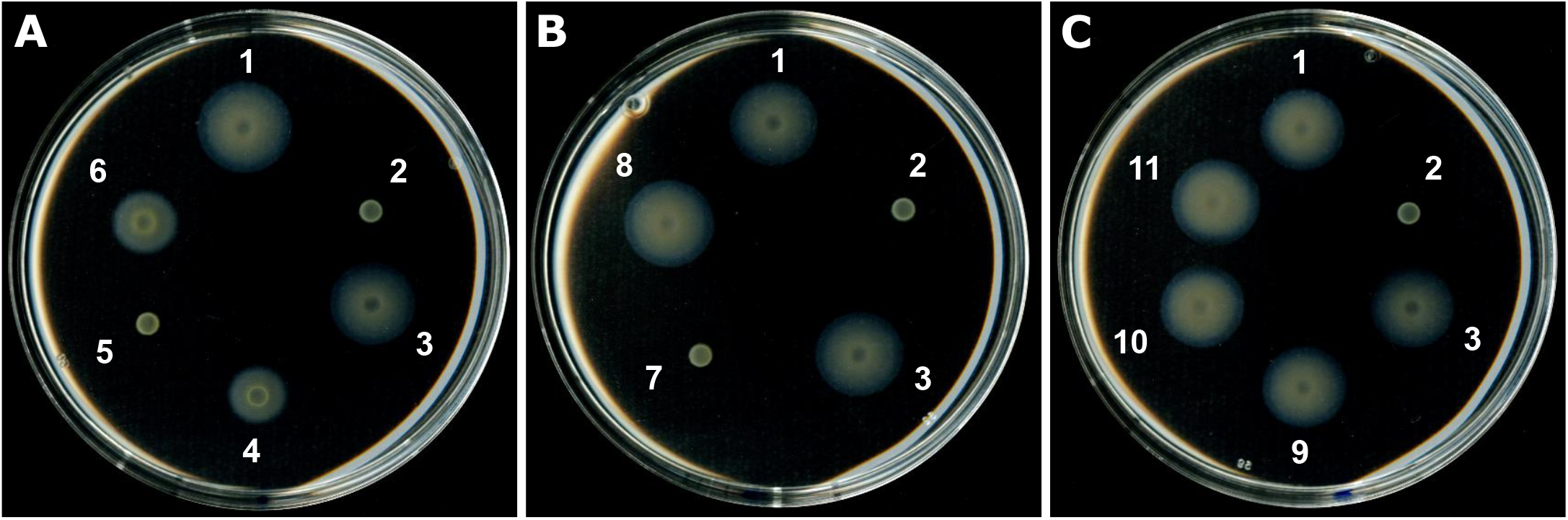
Swimming motility assessment in type IV pilus and flagellum mutants in *Xcc*. To confirm the loss of flagellar motility in the mutants tested in patho-assays, we assayed their ability to move in 0.3% agar swimming plates. 2 µl of bacterial suspensions adjusted to 10^8^ cfu/ml were spotted on swimming plates and pictures were taken after 48 hours of incubation at 28°C. Deletion mutants affected in key components of the type IV pilus (A) and flagellum (B) display reduced and abolished swimming motility respectively. (C) No change in motility was however observed with the Δ*hrpG* and *hrpG*∗ strains. Strains: 1. WT; 2. Δ*fliC*; 3. Δ*fliC::fliC*; 4. Δ*pilA* Δ*pilE*; 5. Δ*pilA* Δ*pilE* Δ*fliC*; 6. Δ*pilA* Δ*pilE* Δf*liC::fliC* 7. Δ*fliQ*; 8. Δf*liQ::fliQ*; 9. Δ*hrpG*; 10. Δ*hrpG::hrpG*; 11. *hrpG*∗

**Figure S4.**
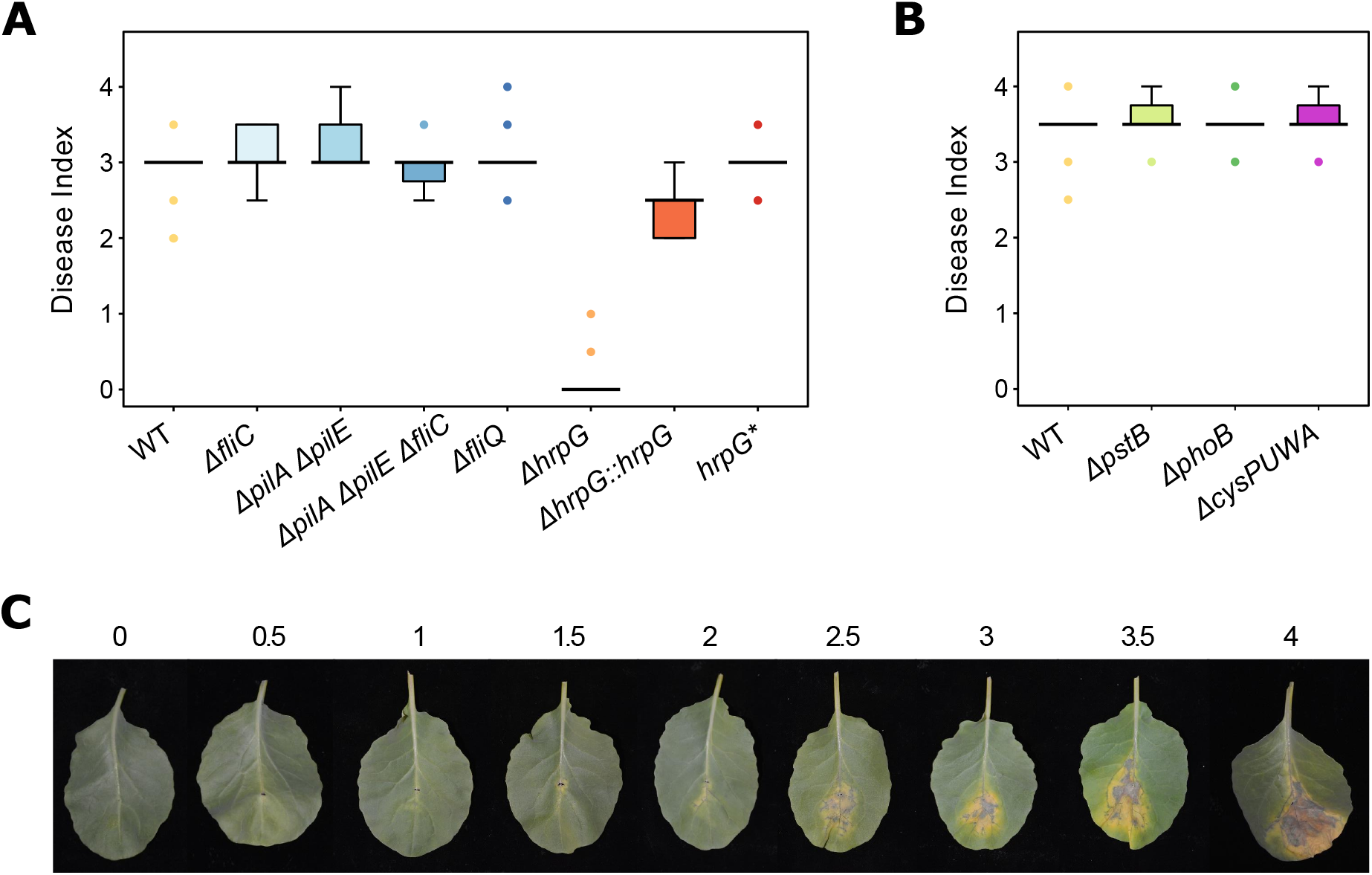
Evaluation of disease symptoms severity during infection of cauliflower leaves by *Xcc* mutants. Severity of the symptoms observed on cauliflower leaves 10 days after inoculation by the *Xcc* 8004::GUS-GFP WT strain or various genetic mutants in (A) motility and T3S regulation or (B) phosphate and sulfate metabolism. The inoculation is performed by piercing the main vein of the second leaf of 4-weeks old cauliflower plants with a needle dipped in a bacterial suspension adjusted to 10^8^ cfu/mL. Boxplots represent the results obtained in three independent biological replicates comprising 5 plants per strain each. (C) Disease Index scale designed in this study to score the severity of the symptoms caused by *Xcc* after inoculation into the main vein of the cauliflower leaf. 0 : No symptoms; 0.5 : Mesophyll discoloration; 1 : Mesophyll discoloration & necrosis along the veins; 1.5 : Vein necrosis & yellow chlorosis at the inoculation point; 2 : Vein necrosis & yellow chlorosis extended to the mesophyll; 2.5 : Extended chlorosis & multiple necrosis spots in the mesophyll away from the inoculation point; 3 : Large chlorosis covering all the diseased area; 3.5 : Large necrosis & large chlorosis area reaching the leaf margin; 4 : Necrosis covering all the diseased area.

**Figure S6.**
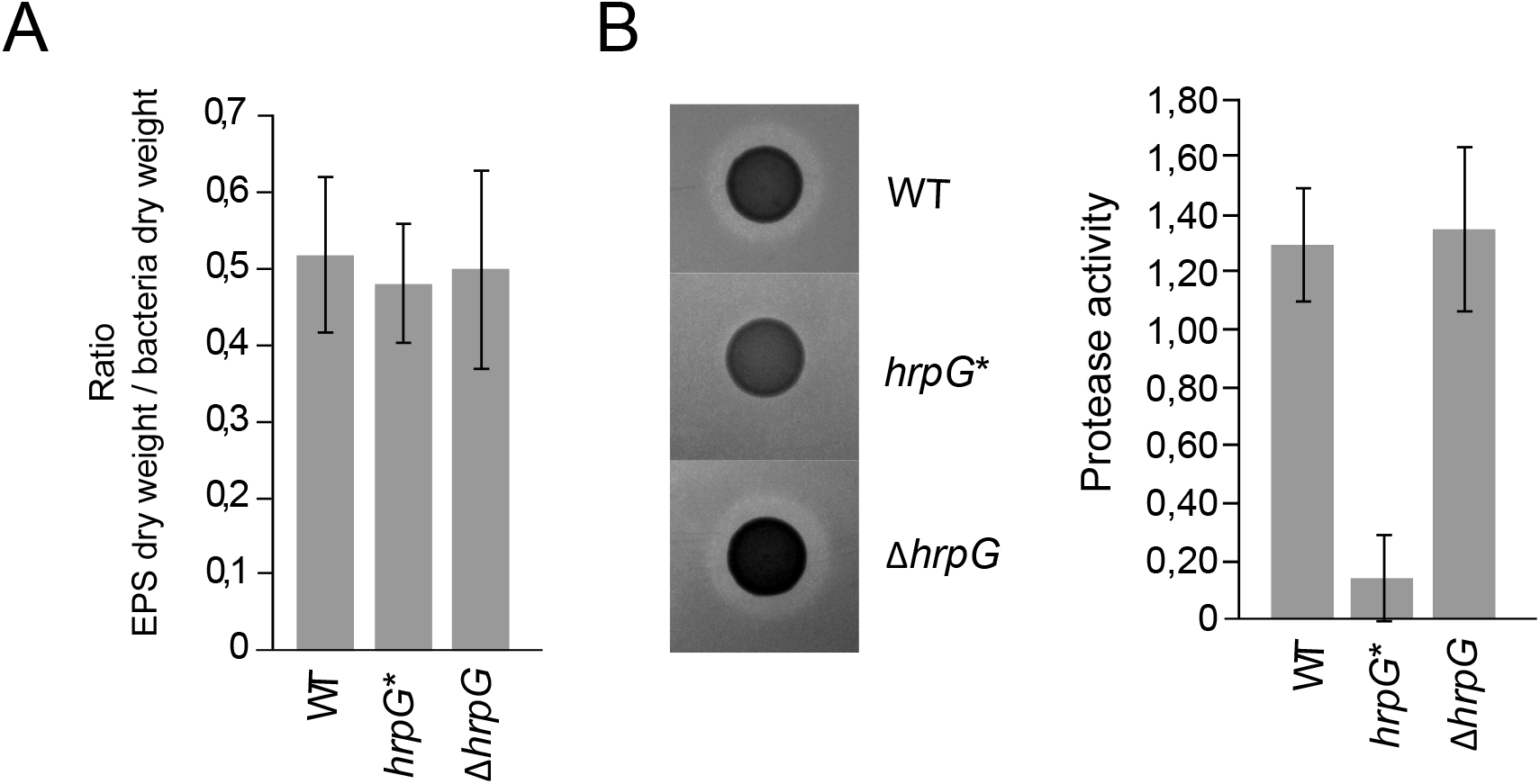
Phenotypic characterization of *hrpG* mutants. (A) EPS production assays were performed on 24 h growth of either 8004::*GUS-GFP* strain (WT), *hrpG*∗ and Δ*hrpG* strains in rich MOKA medium. Production was normalized on dry weight bacteria. Histogram represents the average of results obtained for three independent replicates. (B) On the left, representative pictures of protease activity assay on plate are shown for each tested strain. The histogram represents the average of protease activity measured, as described in the Supplemental Material and Methods section, for three independent replicates.

**Figure S7.**
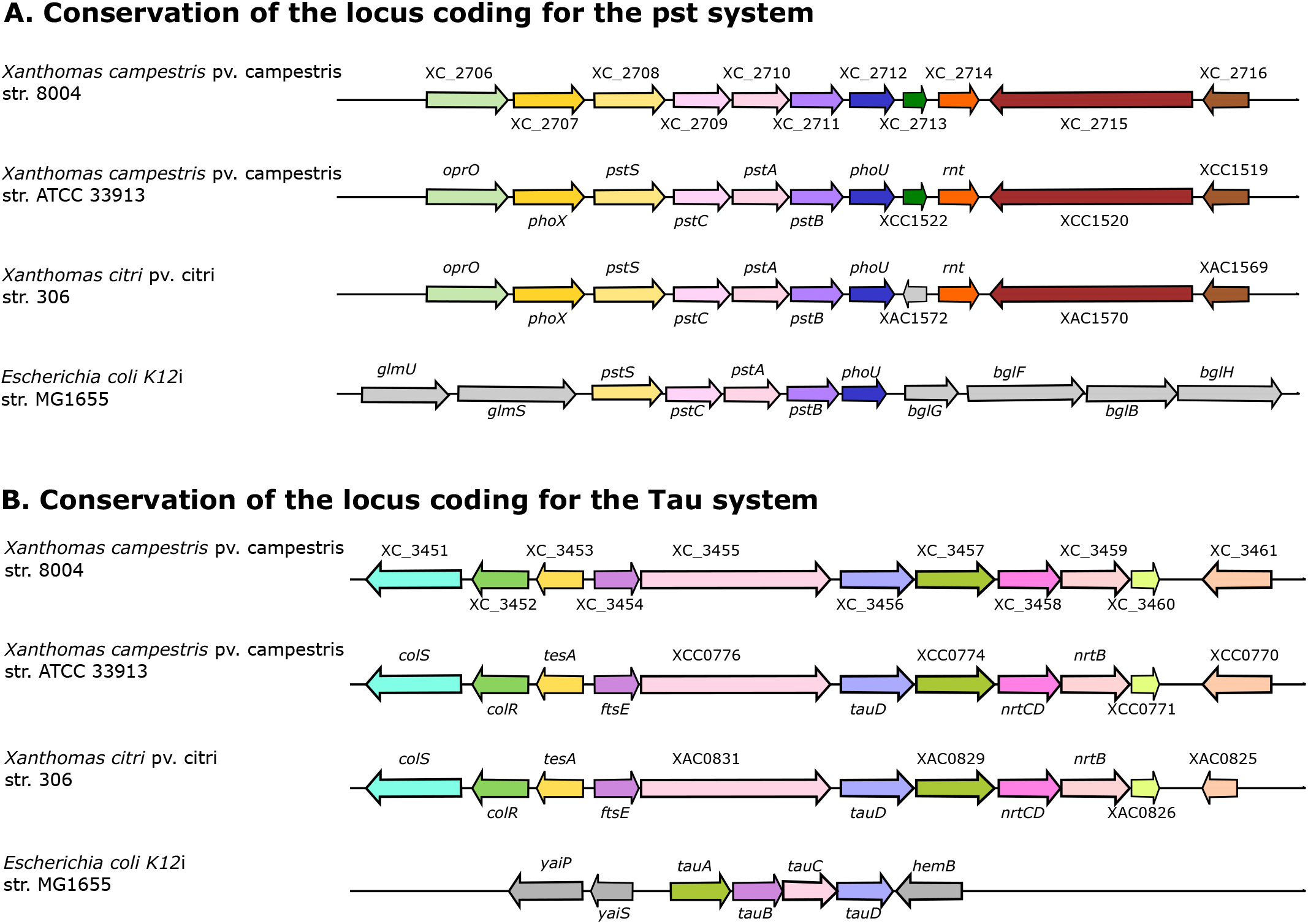
Synteny of genes involved in two transport systems in *Xcc*. Alignment was performed based on protein Blast using the SyntTax tool (https://archaea.i2bc.paris-saclay.fr/synttax/). (A) Comparison of genetic organization of the locus involved in the pst system between *Xcc* strain 8004, *Xcc* strain ATTC33913, *Xac* strain 306 et *E. coli* K12 strain MG1655 using XC_3456 as reference and “Best match” and a minimal threshold normalized BLAST bit score set at 30% as parameters. (B) Comparison of genetic organization of the locus involved in Tau system between *Xcc* strain 8004, *Xcc* strain ATTC33913, *Xac* strain 306 et *E. coli* K12 strain MG1655 using XC_2711 as reference and “Best match” and a minimal threshold normalized BLAST bit score set at 30% as parameters.

## Supplemental Material and Methods

### Bacterial pathogenicity assays

Pathogenicity of *Xcc* strains was tested by wound inoculation of the second true leaf of cauliflower plants by wounding the main vein with a needle dipped in an *Xcc* suspension at 10^8^ cfu/mL. Symptoms were evaluated according to a disease index scale at 7 and 10 dpi (Figure S5). Experiments were performed on five plants per condition and in three independent biological replicates. Significance of differences observed bacterial pathogenicity assays was assessed using the non-parametric Kruskal-Wallis test with **α** = 0.05.

### RNA sequencing procedure

Oriented sequencing was carried out on RNA extracted from *in vitro*-grown *Xcc* by Fasteris SA (Geneva, Switzerland) as described (1). The Small RNA Sequencing Alternative v1.5 Protocol (Illumina) was used for the small RNA fraction, starting with ca. 500 ng RNAs that were treated with tobacco acid pyrophosphatase to remove triphosphate at 5’ transcript ends and purified on acrylamide gel before and after the adaptor ligation step. For large RNAs, a fragmentation step by zinc during 8 min was included before the Illumina procedure. The insert size was 20–120 nt for short RNA libraries and 50–120 nt for long RNA libraries. Libraries were sequenced either in single end (small RNA fraction) or in paired end (Large RNA fraction) on an Illumina HiSeq 2000 platform. Raw sequence data were submitted to the Sequence Read Archive (SRA) database (Accession SRP280320).

For samples prepared from infected tissues, oriented paired‐end RNA sequencing was carried out on the GeT-PlaGe platform (Genotoul, Toulouse, France) using an Illumina HiSeq 3000 platform. Oriented RNA libraries were prepared with TrueSep™ Stranded mRNA Sample Prep Kit (Illumina®). Subsequent steps of sequencing were performed according to the manufacturer instructions (RNA fragmentation, ADNc synthesis, 3’ adenylation). During the fragmentation step, RNAs were cut in fragments of 120 to 210 nt. Raw sequence data were submitted to the Sequence Read Archive (SRA) database (Accession SRP280329).

### Motility assays

*Xcc* cells grown overnight in MOKA were washed by centrifugation (4000 x g, 5 min) and resuspended in sterile water. 2-µl bacterial suspensions adjusted at 10^8^ cfu/mL were spotted on swimming plates (0.03% Bacto peptone, 0.03% yeast extract, 0.3% agar, 2) and incubated 48 h at 28°C. White halos expanding from the colonies indicate the strains’ ability to perform flagellar motility.

### Protease activity assay on milk plates

10 % skimmed milk stock solution was first autoclave for 10 min and used to pour 0.5 % skimmed milk MOKA plates supplemented with 30µg/mL pimaricin. Each plate contains 15 mL of medium. Extracellular protease activity of *Xcc* strains was tested by spotting 5 μL of an overnight culture adjusted to 4.10^8^ cfu/mL. Plates were incubated at 28 °C and imaged 24 hour post inoculation. Diameters of colonies and halos of degradation were measured. Protease activity was calculated as followed: PrA = [π(r^halo^)^2^-π(r^colo^)^2^]/[π(r^colo^)^2^] with “r” corresponding to the radius.

### Measurement of exopolysaccharide production

Extracellular EPS production was measured as described (3). Briefly, an overnight growth of each strain in MOKA was used to inoculate 20 mL of MOKA supplemented with 50µg/mL of rifampicin at 2.10^7^ cfu/mL. After 24 hours of growth 12 mL of culture was centrifuged 15 min at 6500 g. EPS present in supernatant were ethanol precipitated. Then, bacterial and EPS pellets were dried for 8 hours at 65°C before being weighed. EPS production corresponds to the ratio of dry weight EPS on dry weight bacterial cells.

### Growth measurements *in vitro*

*In vitro* growth curves were generated using a FLUOStar Omega apparatus (BMG Labtech, Offenburg, Germany) using 96-well flat-bottom microtiter plates (Greiner) with 200 µL of bacterial suspensions. After an overnight preculture in MOKA rich medium, cells were harvested by centrifugation at 9500 g for 4 minutes, washed and resuspended in MME minimal medium (K2HPO4 10.5 g/L, KH2PO4 4.5 g/L, (NH4)2SO4 1 g/L MgSO4 0.12g/L, casamino acids 0.15 g/L) (4). Bacterial suspensions inoculated at an optical density at 600 nm (OD600) of 0.15 were prepared in MME and MOKA media. For each experiment, four replicates coming from two independent precultures were performed. The microplates were shaken continuously at 700 rpm using the linear-shaking mode. Each experiment was repeated three times and a representative experiment was shown.

## Notes

### Competing Interest Statement

The authors have declared no competing interest.

