## Supplementary material for "*Xanthomonas* transcriptome inside cauliflower hydathodes reveals bacterial virulence strategies and physiological adaptation at early infection stages": Table S1

**Table S1: List of strains and vectors used in this study**

| Strain or Plasmid | Relevant characteristics | Reference or source |
|---|---|---|
| ***Xanthomonas campestris* pv. *campestris*** | | |
| 8004 | Wild-type, spontaneous Rifampicin-resistant mutant of NCPPB 1145; $Rif^R$ | {Daniels, 1984 #6958} |
| 8004::*hrpG** | *hrpG*$_{E44K}$ mutant derivative, constitutive *hrp* gene expression; $Rif^R$ | {Guy, 2013 #7382} |
| 8004::*GUS-GFP* | Wild-type, constitutive *GUS* and *GFP* expression; $Rif^R$ | {Cerutti, 2017 #7654} |
| 8004::*GUS-GFP ΔhrpG* | *hrpG* (*XC_3077*) deletion mutant, constitutive *GUS* and *GFP* expression; $Rif^R$ | This study |
| 8004::*GUS-GFP ΔhrpG::hrpG* | *hrpG* complemented mutant, constitutive *GUS* and *GFP* expression; $Rif^R$ | This study |
| 8004::*GUS-GFP hrpG** | *hrpG*$_{E44K}$ mutant derivative, constitutive *hrp* gene expression, constitutive *GUS* and *GFP* expression; $Rif^R$ | This study |
| 8004::*GUS-GFP ΔfliC* | *fliC* (*XC_2245*) deletion mutant, constitutive *GUS* and *GFP* expression; $Rif^R$ | This study |
| 8004::*GUS-GFP ΔfliC::fliC* | *fliC* complemented mutant, constitutive *GUS* and *GFP* expression; $Rif^R$ | This study |
| 8004::*GUS-GFP ΔpilAΔpilE* | *pilA* (*XC_1058*) *pilE* (*XC_1626*) double deletion mutant, constitutive *GUS* and *GFP* expression; $Rif^R$ | This study |
| 8004::*GUS-GFP ΔpilAΔpilEΔfliC* | *pilA* (*XC_1058*) *pilE* (*XC_1626*) *fliC* (*XC_2245*) triple deletion mutant, constitutive *GUS* and *GFP* expression; $Rif^R$ | This study |
| 8004::*GUS-GFP ΔpilAΔpilEΔfliC::fliC* | *pilA pilE fliC* triple deletion mutant complemented with *fliC*, constitutive *GUS* and *GFP* expression; $Rif^R$ | This study |
| 8004::*GUS-GFP ΔfliQ* | *fliQ* (*XC_2272*) deletion mutant, constitutive *GUS* and *GFP* expression; $Rif^R$ | This study |
| 8004::*GUS-GFP ΔfliQ::fliQ* | *fliQ* complemented mutant, constitutive *GUS* and *GFP* expression; $Rif^R$ | This study |
| 8004::*GUS-GFP ΔpstB* | *pstB* (*XC_2711*) deletion mutant, constitutive *GUS* and *GFP* expression; $Rif^R$ | This study |
| 8004::*GUS-GFP ΔpstB::pstB* | *pstB* complemented mutant, constitutive *GUS* and *GFP* expression; $Rif^R$ | This study |
| 8004::*GUS-GFP ΔphoB* | *phoB* (*XC_3272*) deletion mutant, constitutive *GUS* and *GFP* expression; $Rif^R$ | This study |
| 8004::*GUS-GFP ΔphoB::phoB* | *phoB* complemented mutant, constitutive *GUS* and *GFP* expression; $Rif^R$ | This study |
| ***E. coli*** | | |

| | | |
|---|---|---|
| TG1 | *supE thi-1 Δ(lac-proAB) Δ(mcrB-hsdSM)5 (*$rk^-$ $mk^-$*) [F′ traD36 proAB lacI*$^q$ *ZΔM15]* | Stratagene |
| DH5$\alpha$ | $F^-$ *recA lacZ DM15* | Bethesda Research Laboratory |
| **Plasmids** | | |
| pRK2073 | Helper plasmid for triparental mating; $Spec^R$; $Strep^R$ | {Leong, 1982 #7146} |
| pK18mobSacB | Suicide vector in *Xcc* used for genome edition; $Kan^R$ | {Schafer, 1994 #6023} |
| pK18_XC3077_AmAv | *XC_3077* deletion construct; pK18mobSacB derivative; $Kan^R$ | This study |
| pK18_XC2245_AmAv | *XC_2245* deletion construct; pK18mobSacB derivative; $Kan^R$ | This study |
| pK18_XC1058_AmAv | *XC_1058* deletion construct; pK18mobSacB derivative; $Kan^R$ | This study |
| pK18_XC1626_AmAv | *XC_1626* deletion construct; pK18mobSacB derivative; $Kan^R$ | This study |
| pK18_XC2272_AmAv | *XC_2272* deletion construct; pK18mobSacB derivative; $Kan^R$ | This study |
| pK18_XC2711_AmAv | *XC_2711* deletion construct; pK18mobSacB derivative; $Kan^R$ | This study |
| pK18_XC3272_AmAv | *XC_3272* deletion construct; pK18mobSacB derivative; $Kan^R$ | This study |
| pK18_compR3 | pK18mobSacB derivative use for genomic complementation, pTac promoter; $Kan^R$ | This study |
| pK18_Comp_XC3077 | *XC_3077* overexpression construct; pK18_compR3 derivative; $Kan^R$ | This study |
| pK18_Comp_XC2245_5UTR | *XC_2245* overexpression construct; pK18_compR3 derivative; $Kan^R$ | This study |
| pK18_Comp_XC2272 | *XC_2272* overexpression construct; pK18_compR3 derivative; $Kan^R$ | This study |
| pK18_Comp_XC3272 | *XC_3272* overexpression construct; pK18_compR3 derivative; $Kan^R$ | This study |
| pK18_compR3_RBS | pK18mobSacB derivative use for genomic complementation, pTac promoter with ribosome-binding site added; $Kan^R$ | This study |
| pK18_CompRBS_XC2711 | *XC_2711* overexpression construct; pK18_compR3_RBS derivative; $Kan^R$ | This study |

Rif: Rifampicin; Kan: Kanamycin; Spec : Spectinomicin; Strep: Streptomicin; RBS: Ribosome Binding Site
