## Supplementary material for "*Xanthomonas* transcriptome inside cauliflower hydathodes reveals bacterial virulence strategies and physiological adaptation at early infection stages": Table S2

**Table S2: Sequences of oligonucleotides used for construction of deletion and complementation plasmids**

| Locus name | Position | Direction | Name | Sequence (5’-3’) |
|---|---|---|---|---|
| **Plasmids construction** | | | | |
| *XC_1301* | Right | Fw | pK18_Am1301 | *TGTAAAACGACGGCCAGTGCCAAGCT*CAATCGCACGTGATGGAGCGCTGG |
| | Left | Rev | MCS_Am1301 | *TATGATGTCGGCGCAAAAACTGCAG*GGCAAGGCCGTTGTGGCGACTCGGC |
| *XC_1302* | Right | Rev | pK18_Av1302 | *GAGTCGACCTGCAGGCATGCAAGCT*GGCTATGGCGCCTGGGAAATCGGCC |
| | Left | Fw | MCS_Av1302 | *TACGCGCAGCGTGACCGCTACACTT*GCTGTGGTGATCAGCGGGCCGCAGC |
| pTac-T7 MCS region | Right | Fw | Am1301_MCS | AGCGCCGAGTCGCCACAACGGCCTTGCC*CTGCAGTTTTTGCGCCGACATC* |
| | Left | Rev | Av1302_MCS | GCGCTGCGGCCCGCTGATCACCACAGC*AAGTGTAGCGGTCACGCTGCGCG* |
| RBS | Right | Fw | CompR3_RBSup | *CGGATCCGAATTCGAGCTCCGTCGACAAGCTGAATTAATTCGGATCCCTC* |
| | Left | Rev | CompR3 _RBSdw | *CGCGTTTAAACCGCATGCATCGCAAGCTTGGTCGAGATCCTCTAGAGCTC* |
| **Deletion** | | | | |
| *XC_3077* | Left | Fwd | pK18_3077Am | *GTAAAACGACGGCCAGTGCCAAGCT*CGGTCCACCTGTAATTACAAGAGC |
| | | Rev | XC_3077AvAm | CTCAGCAAGCTGCGGTGCGATTGACGTCATTCGCAGCGGCGTCCATCAC |
| | Right | Fwd | XC_3077AmAv | GTGATGGACGCCGCTGCGAATGACGTCAATCGCACCGCAGCTTGCTGAG |
| | | Rev | pK18_3077Av | *GAGTCGACCTGCAGGCATGCAAGCT*GTACGGCGAGAGCCCGAAGCAG |

| | | | | |
|---|---|---|---|---|
| *XC_2245* | Left | Fwd | pK18_2245Am | *GTAAAACGACGGCCAGTGCCAAGCT*CGCTCAAGACCACCTTGTCGTCGAT |
| | | Rev | XC_2245AvAm | GCTGTGACGCTGCCGATTACTGCAGCAGCGACATTACGTTGGTGTTGATTACCTGTG |
| | Right | Fwd | XC_2245AmAv | AATCAACACCAACGTAATGTCGCTGCTGCAGTAATCGGCAGCGTCACAGCAACGC |
| | | Rev | pK18_2245Av | *GAGTCGACCTGCAGGCATGCAAGCT*CCGAGCTGGTGGTGGTGGCGGTAAA |
| *XC_1058* | Left | Fwd | pK18_1058Am | *GTAAAACGACGGCCAGTGCCAAGCT*GAGCATGTTCGCGTTGCGAGCAGG |
| | | Rev | XC_1058AvAm | GAGCTTTGATTACGGAGCCGCCGCTTCGATCAGTGTAAAACCATTTTGC |
| | Right | Fwd | XC_1058AmAv | GCAAAATGGTTTTACACTGATCGAAGCGGCGGCTCCGTAATCAAAGCTC |
| | | Rev | pK18_1058Av | *GAGTCGACCTGCAGGCATGCAAGCT*ACTGAATAGTAGTCCCGGAGCCGTTG |
| *XC_1626* | Left | Fwd | pK18_1626Am | *GTAAAACGACGGCCAGTGCCAAGCT*CAGGATGGTGATCGGACATTGAGCC |
| | | Rev | XC_1626AvAm | GCCAGTCTTTACATCACCAACAGTCACGCCCCCTAACTGCTTGCAAGAAC |
| | Right | Fwd | XC_1626AmAv | GTTCTTGCAAGCAGTTAGGGGGCGTGACTGTTGGTGATGTAAAGACTGGC |
| | | Rev | pK18_1626Av | *GAGTCGACCTGCAGGCATGCAAGCT*CAGACAGCAACTGGTCCGCAGGGTG |
| *XC_2272* | Left | Fwd | pK18_2272Am | *GTAAAACGACGGCCAGTGCCAAGCT*ATGGGGGACCCCGAAGCAGAACAGC |
| | | Rev | XC_2272AvAm | TCCATCAGCGCGTGCCGCTACCCGATAATCCACAAGACGGTGACCAGGCC |
| | Right | Fwd | XC_2272AmAv | TGGCCTGGTCACCGTCTTGTGGATTATCGGGTAGCGGCACGCGCTGATGG |
| | | Rev | pK18_2272Av | *AGAGTCGACCTGCAGGCATGC* |

| | | | | |
|---|---|---|---|---|
| | | | | *AAGCT*TAGCTGTCCACCAGCAACGCGATC |
| *XC_2711* | Left | Fwd | pK18_2711Am | *GTAAAACGACGGCCAGTGCCAAGCT*GCTGTACGTGATGCAGACCG |
| | | Rev | XC_2711AvAm | GCAGGCCTACATCAGCCGAAGCGGCTTGGCGTTTTGGAGATCGTTC |
| | Right | Fwd | XC_2711AmAv | GAACGATCTCCAAAACGCCAAGCCGCTTCGGCTGATGTAGGCCTGC |
| | | Rev | pK18_2711Av | *GAGTCGACCTGCAGGCATGCAAGCT*GGAATTGAGTGCGATCGAGCGCTTG |
| *XC_3272* | Left | Fwd | pK18_3272Am | *GTAAAACGACGGCCAGTGCCAAGCT*GCTTCGATGCGCTGCTGAAGC |
| | | Rev | XC_3272AvAm | GGCATCAGGTGGCCGAGGAGAAACGGTCATCGACGATCAGGATGCG |
| | Right | Fwd | XC_3272AmAv | CGCATCCTGATCGTCGATGACCGTTTCTCCTCGGCCACCTGATGCC |
| | | Rev | pK18_3272Av | *GAGTCGACCTGCAGGCATGCAAGCT*GCCTTGTTGAACCACTGCAC |
| **Complementation** | | | | |
| *XC_3077* | Left | Fwd | pK18_CompR3_XC3077_Fwd | *CGAATTCGAGCTCCGTCGACAAGCT*CTGTTGTGCTCAGTTCCATGGTCTG |
| | Right | Rev | pK18_CompR3_XC3077_Rev | *GTTTAAACCGCATGCATCGCAAGCT*TCAGCAAGCTGCGGTGCGATTGACC |
| *XC_2245* | Left | Fwd | pK18_Comp_XC2245_5UTR_Fwd | *CGAATTCGAGCTCCGTCGACAAGCT*CTAAAGGTTCCCAGAGCATCG |
| | Right | Rev | pK18_Comp_XC2245_out_Rev | *GTTTAAACCGCATGCATCGCAAGCT*GCTCTAGCGTTGCTGTGACG |
| *XC_2272* | Left | Fwd | pK18_CompR3_XC2272_Fwd | *CGAATTCGAGCTCCGTCGACAAGCT*GCACCGCACTTGCATCAGCACTCC |
| | Right | Rev | pK18_CompR3_XC2272_Rev | *GTTTAAACCGCATGCATCGCAAGCT*CTACCCGATCAGATGCGGAATGC |

| | | | | |
|---|---|---|---|---|
| *XC_2711* | Left | Fwd | pK18_CompRBS_XC2711_Fwd | *GCTCTAGAGGATCTCGACCAAGCT*ATGACTGACCTTTCTTTCCGGACG |
| | Right | Rev | pK18_CompRBS_XC2711_Rev | *CGTTTAAACCGCATGCATCGCAAGCT*TCAGCCGAAGCGGCCGGTGATG |
| XC_3272 | Left | Fwd | pK18_CompR3_XC3272_Fwd | *CGAATTCGAGCTCCGTCGACAAGCT*CGTAGTCTACTGGCGACCATCC |
| | Right | Rev | pK18_CompR3_XC3272_Rev | *GTTTAAACCGCATGCATCGCAAGCT*TCAGGTGGCCGAGGAGAAACGGTAG |

Sequences corresponding to restriction sites are underlined and sequences non homologous to *Xcc* are represented in italics.
