## Supplementary material for "*Xanthomonas* transcriptome inside cauliflower hydathodes reveals bacterial virulence strategies and physiological adaptation at early infection stages": Table S3

### Table S3: Sequences of oligonucleotides used for oligocapture of plant RNAs

| Name | Sequence (5’-3’)[a] |
|---|---|
| **Nuclear ribosomal RNA** | |
| 28S_2844_LNA | CTC{C}CGA{C}AATTTCAA{G}CA{C}TCT |
| 28S_3090_LNA | GT{G}TTT{C}AA{G}ACGGGTCGAA{T}GG |
| 28S_3345_LNA | C{C}TATACCCAAGT{C}AGA{C}GAA{C}G |
| 28S_3660_LNA | T{C}TTAAT{C}GAC{C}AACACC{C}TTTG |
| 28S_3889_LNA | G{C}TACTA{C}CAC{C}AAGATCTG{C}AC |
| 28S_4534_LNA | CCTA{C}ATTGTT{C}CATC{G}AC{C}AGA |
| 28S_4803_LNA | GGA{C}CATC{G}CAATG{C}TTTGT{T}TT |
| 28S_5082_LNA | GT{C}ATTTCA{C}AAAGTCG{G}ACT{A}G |
| 28S_5399_LNA | GA{T}AGG{C}CACG{C}TTTCACGG{T}TC |
| 28S_5772_LNA | CG{G}ATTCTGA{C}TTAGA{G}GCG{T}TC |
| 28S_5693_LNA | GA{C}CAATTGT{G}CGAA{T}CAACG{G}T |
| 28S_6158_LNA | AA{G}TTGG{G}AATT{C}GTTAAG{G}AGC |
| 28S_2474_LNA | TGA{T}ATGCT{T}AAA{C}TCAGC{G}GGT |
| 28S_2707_LNA | CCAAA{C}AACCCGA{C}TC{G}TAGA{C}A |
| 28S_4147_LNA | TT{T}CCAG{G}GTGGG{C}AGG{C}TGTTA |
| 18S_70_LNA | GTT{C}ATA{C}TTA{C}ACATGCATG{G}C |
| 18S_174_LNA | GCA{C}GTATT{A}GCT{C}TAGAA{T}TAC |
| 18S_383_LNA | AT{C}GAA{C}CCTAATTCT{C}CGT{C}AC |
| 18S_1792_LNA | CG{G}AAA{C}CTTGTTACGA{C}TT{C}TC |
| 18S_1677_LNA | ATT{C}AATCG{G}TAG{G}AGCGA{C}GGG |

| | |
|---|---|
| 18S_559_LNA | TG{C}CCT{C}CAATGGAT{C}CTC{G}TTA |
| 18S_258_LNA | AT{C}ATGAA{T}CATCAG{A}GCAA{C}GG |
| 18S_1147_LNA | TTC{C}TTTAAGTT{T}CAG{C}CTTG{C}G |
| 18S_1412_LNA | GG{C}CATAGTC{C}CTCTAA{G}AA{G}CC |
| 28S_843_LNA | GCCAA{C}ACAA{T}AGGA{T}CGAA{A}TC |
| 5,8S_2159_LNA | ACA{C}CAA{G}TATCGCA{T}TT{C}GCT |
| **Choroplastic ribosomal RNA** | |
| Bo0151s070_23S_10_LNA | A{C}AGCTT{C}GGCA{G}ATCG{C}TTAGC |
| Bo01051s070_23S_256_LNA | CAAG{G}GGTA{G}TA{C}AGGAATA{T}TC |
| Bo01051s070_23S_832_LNA | AGA{C}AGTG{C}CCA{G}ATCGT{T}ACGC |
| Bo01051s070_23S_974_LNA | CCTT{C}TTCG{C}CTT{C}CA{C}CTAAGC |
| **Other plant RNA** | |
| BRARA_CHL_psbA_43_LNA | GTTAT{C}CAGTTA{C}AGAA{G}C{G}ACC |
| BRARA_CHL_psbA_199_LNA | GTTT{C}TGGAT{C}T{C}TTCTTTA{C}GG |
| BRARA_CHL_psbA_506_LNA | GATT{C}CTA{G}AGG{C}ATACCAT{C}AG |
| BRARA_CHL_psbA_745_LNA | CCA{A}AATAA{C}CGTGA{G}CAG{C}TAC |
| BRARA_CHL_psbA_829_LNA | CAA{A}TACCTA{C}TAC{C}GGC{C}AAGC |
| BRARA_CHL_psbA_932_LNA | CA{G}CC{C}AAGTATT{A}ATAACA{C}GTC |
| BRARA_CHL_rbcL_4_LNA | G{C}TTTAGT{C}TCT{G}TTTGTGGT{G}A |
| BRARA_CHL_rbcL_188_LNA | C{C}ACA{C}AGTTGTC{C}ATGTA{C}CAG |
| BRARA_CHL_rbcL_731_LNA | CA{T}CATTT{C}TTCG{C}ATGTA{C}CCG |
| BRARA_CHL_rbcL_1032_LNA | CAT{C}GCG{C}AG{T}AAATCAA{C}AAAG |
| BRARA_CHL_rbcL_1395_LNA | CGAT{G}GTTGG{G}AA{G}TTAAAT{G}TG |

| | |
|---|---|
| Bo01051s080_29_LNA | GTAA{C}CCAT{G}CTAT{A}CTC{C}CAGG |
| Bo01051s080_127_LNA | GGTTCAA{G}AACG{C}AAG{G}TGT{C}CC |
| Bo01051s080_517_LNA | CTA{G}AAGCAG{C}CA{C}CCTTG{A}AAG |
| Bo01051s080_709_LNA | CGGTA{C}CAAAT{C}GAG{G}CAA{A}CTC |
| Bo01051s080_812_LNA | GAAG{C}TTAT{C}CCCCA{C}CGT{C}TCA |

[a] Bases under brackets refer to locked nucleic acid (LNA)-modified bases
