## Supplementary material for "*Xanthomonas* transcriptome inside cauliflower hydathodes reveals bacterial virulence strategies and physiological adaptation at early infection stages": Table S8

**Table S8: Properties of RNAseq libraries from *Xcc* strain 8004 wild-type and derivatives grown *in vitro* or harvested from cauliflower hydathodes**

| RNA size | Strain | Condition | Biological replicates | Number of raw reads | Uniquely mapped reads *Xcc* & *Brassica* (%)[a] | Hits *Brassica* genes (%)[b] | Hits *Xcc* genes (%)[c] | Accession[d] |
|---|---|---|---|---|---|---|---|---|
| **Long RNAs fraction** | 8004 | MOKA | 3 | 22 503 680 | 93.58 | - | 9472 | SRR12603737 |
| | | | | 31 781 601 | 88.67 | - | 9449 | SRR12603738, SRR12603725 |
| | | | | 29 317 932 | 88.54 | - | 9421 | SRR12603739, SRR12603726 |
| **Small RNAs fraction** | | | 3 | 14 141 422 | - | - | - | SRR12603734 |
| | | | | 13 443 960 | - | - | - | SRR12603735 |
| | | | | 13 423 070 | - | - | - | SRR12603736 |
| **Long RNAs fraction** | 8004::*hrpG** | MOKA | 3 | 21 746 450 | | - | 9358 | SRR12603727 |
| | | | | 14 282 665 | 94.84 | - | 9411 | SRR12603728 |
| | | | | 15 932 094 | 95.08 | - | 9457 | SRR12603729 |
| **Small RNAs fraction** | | | 4 | 10 314 675 | - | - | - | SRR12603720 |
| | | | | 11 898 842 | - | - | - | SRR12603721 |
| | | | | 10 794 476 | - | - | - | SRR12603722 |
| | | | | 14 113 815 | - | - | - | SRR12603733 |
| | 8004 | Cauliflower hydathodes 4 hpi | 3 | 158 422 507 | 33.42 | 6245 | 1332 | SRR12603813, SRR12603814, SRR12603815 |
| | | | | 150 271 178 | 23.49 | 6621 | 1063 | SRR12603810, SRR12603811, SRR12603812 |
| | | | | 173 591 985 | 41.15 | 6794 | 1012 | SRR12603804, SRR12603805, SRR12603806 |
| | 8004 | Cauliflower hydathodes 72 hpi | 3 | 164 579 058 | 38.59 | 5519 | 1640 | SRR12603816, SRR12603817, SRR12603818 |
| | | | | 163 894 181 | 38.47 | 5256 | 2727 | SRR12603801, SRR12603802, SRR12603803 |
| | | | | 143 337 188 | 27.63 | 73.48 | 1.61 | SRR12603807. SRR12603808, SRR12603809 |

[a] Percentage of raw reads mapped uniquely to the sequences of *Xcc* strain 8004 chromosome (Genbank accession number CP000050.1), *Brassica oleracea* nuclear and mitochondrial genomes (Accession GCA_000695525.1, NC_016118.1) and *Brassica rapa* chloroplastic genome (BRARA_CHL, accession NC_040849.1).
[b] Percentage of unique hits to the genes of *Brassica oleracea* nuclear and mitochondrial genomes and *Brassica rapa* chloroplastic genome.
[c] Percentage of unique hits to the genes of *Xcc* strain 8004.
[d] Sequence Read Archive accession number
